# A 20-years comparative study of domoic acid depuration in king scallops, *Pecten maximus*, across French provinces

**DOI:** 10.1101/2025.05.08.652798

**Authors:** Eline Le Moan, Amélie Derrienb, Caroline Fabioux, Fred Jean, Malwenn Lassudrie, Aourégan Terre-Terrillon, Hélène Hégaret, Jonathan Flye-Sainte-Marie

## Abstract

Harmful Algal Blooms (HABs) can lead to fishery closures when toxin levels in commercial species exceed regulatory thresholds. Domoic acid (DA), the neurotoxin causing Amnesic Shellfish Poisoning (ASP), is particularly persistent in king scallops (*Pecten maximus*), a commercially valuable species in France. This species is known for its slow depuration rate compared to other species. As a result, anticipating DA dynamics is therefore a key to managing fishery openings in autumn, especially following spring contamination events. This study used 20 years of data from the French phycotoxin *in-situ* monitoring programme (REPHYTOX) to identify contamination events along the French Atlantic and English Channel coastlines. Depuration rates were estimated for 104 events; however, no correlation was found between depuration rate and province, time period, initial DA concentration or environmental conditions. Consequently, a median depuration rate was defined and applied in a widely used exponential decay model. In response to professionals’ needs, we developed a user-friendly predictive tool that estimates DA concentrations in king scallops based on sampling in spring or summer. We recommend performing DA quantification in scallops whenever DA is detected in other shellfish at the same location, and running the predictive model to anticipate DA content at the opening of the fishery season in autumn. This tool will help fishery managers to anticipate bans, to avoid unnecessary licence purchases, or to shift to alternative species. While developed using French data, the methodology is adaptable to other regions, with appropriate adjustments to reflect local ecological, regulatory and fishery contexts.

## 1. Introduction

Harmful Algal Blooms (HABs) are defined as the natural proliferation of microalgae that produce toxic compounds which can be responsible for human poisoning syndromes, including Amnesic Shellfish Poisoning (ASP), Paralytic Shellfish Poisoning (PSP), and Diarrhetic Shellfish Poisoning (DSP) (Sellner et al., 2003). In humans, ASP is characterised by symptoms such as nausea, vomiting, memory loss and confusion, which can potentially evolve into coma and death (Pulido, 2008). This syndrome is caused by the neurotoxin domoic acid (DA), produced by several microalgal species, particularly pelagic diatoms of the ubiquitous genus *Pseudo-nitzschia*. A total of 62 species have been identified (Guiry and Guiry, 2021) of which 29 are toxigenic (Lundholm et al., 2009). The first case of human intoxication from domoic acid was reported in 1987 in Canada, leading to three deaths and 153 cases of poisoning Bates et al. (1988). This marked the first instance of domoic acid being identified as the cause of ASP symptoms and deaths, following the consumption of contaminated mussels (Wright et al., 1989; Bates et al., 1988). Subsequently, this toxin has been monitored in commercially exploited shellfish species to prevent intoxication in numerous countries, including France (https://bulletinrephytox.fr/accueil, REPHYTOX), Ireland (Marine Institute: https://webapps.marine.ie/HABs/), the United Kingdom (Food Standards Agency: https://www.food.gov.uk/business-guidance/biotoxin-and-phytoplankton-monitoring) and Portugal (Instituto Português do Mar e da Atmosfera: https://www.ipma.pt/pt/bivalves/index.jsp). An international regulatory threshold of 20 *mg DA kg*^−1^ total flesh weight (Commission, 2002) has been established, above which the exploitation is prohibited to prevent human poisoning.

Species that feed on toxin-producing microalgae are particularly exposed to HABs and can act as vectors for toxin transfers within the food web. This is particularly true for filter-feeding species such as bivalves. They can accumulate toxins in their tissues through feeding, making them unsuitable for human consumption (MacDonald et al., 2016). The main strategy to prevent intoxication and associated health complications is to impose sale bans by closing fisheries and/or suspending the commercialisation of aquaculture and fishery products. However, bivalves are of high economic value, accounting for 1.4 million tonnes from captures (FAO, 2025b) and 17 million tonnes from aquaculture globally in 2022 (FAO, 2025a). Pectinid species, including king scallops (*Pecten maximus* Linnaeus, 1758), queen scallops (*Aequipecten opercularis* Linnaeus, 1758), and variegated scallops (*Mimach-lamys varia* Linnaeus, 1758), accounted for 710 000 tonnes of capture and 2 million tonnes in aquaculture, thus accounting for 50% of total bivalve fisheries and 11% of global bivalve aquaculture (FAO, 2025a,b). The aquaculture of pectinid species generated USD 5 billion in 2022. Consequently, the closure of aquaculture and fishing activities result in significant socio-economic losses. Several European countries are affected by HABs and particularly ASP events. In Galicia, for example, king scallop harvesting is permanently banned. However, professionals can request for domoic acid controls in tissues. If concentrations range between 20 and 250 mg kg-1, shucking is allowed under certain conditions (European Commission, 2002). Shucking removes the digestive gland and the soft tissues - where toxin levels are highest - leaving only the muscle, with or without gonad, to ensure levels stay below the 20 *mg DA kg*^−1^ limit for sale. Consequently, the exploitation is sporadic, and controls are only made on demand. In Ireland, according to the bulletins issued by the Marine Institute (https://www.marine.ie/site-area/data-services/interactive-maps/weekly-hab-bulletin), king scallops are regularly contaminated by domoic acid. However, the sale of adductor muscle only, with or without gonads, or of the entire organism can be authorised depending on domoic acid concentration.

Events of ASP were first observed along the French Atlantic coastlines in 2004 and currently, all French coasts (Mediterranean, Atlantic and English Channel) face ASP, DSP and PSP events (Belin et al., 2021). Data come from REPHY (REPHY - French Observation and Monitoring program for phytoplankton and hydrology in coastal waters, 2023) and REPHYTOX (REPHYTOX - French Monitoring program for phycotoxins in marine organisms, 2023) programmes which monitor the biomass, abundance and composition of marine phytoplankton communities as well as regulated phycotox-ins in molluscs since 1984. King scallops were the first species sold in fish auctions in terms of volumes in 2021 and 2022 (FranceAgriMer, 2022). REPHYTOX, tied to REPHY, triggers shellfish sampling when toxin-producing phytoplankton are present. Offshore exploited species (*e.g*., king scallops, clams, common European bittersweet) are sampled biweekly during fishing if toxin levels remain below legal limits.

French areas most affected by all types of HABs over the 2000-2018 period were the Bay of Seine in Normandy and the western and southern coasts of Brittany (Belin et al., 2021). This study found that 17 to 19 out of 19 years were affected by DSP for both regions and between 5 and 8 years for ASP in the Bay of Seine and between 9 and 16 years for western and southern Brittany. Incidence of ASP along the French coastlines has increased over the past two decades especially linked to increase in toxin monitoring with the obligation to monitor scallops. Scallop fishing is still practised in France; however, the presence of toxins responsible for DSP, PSP and ASP has been documented at different frequencies depending on the regions, which can result in fishery closures. Studies highlight that the economic impact of HABs depends on the duration and the geographic extent of closures (Holland and Leonard, 2020; Chenouf et al., 2023). Guillotreau et al. (2021) emphasise that the professionals’ ability to anticipate HAB events is a key factor in the success of adaptive strategies. Moreover, the impact of HAB-related closures on scallop fishing activities depends on fleet mobility, as described in Chenouf et al. (2023). Activities with a limited adaptive capacity, such as relocating from an affected zone to a non-affected zone, are especially vulnerable to severe impacts. This strategy’s feasibility depends on the size of the fishing area. For example, in the small zone of the Bay of Brest (France), where king scallops are the primary pectinid species being fished, ASP events prevent fishermen from relocating their activities to an alternative zone, unlike in the larger area of Bay of Seine (France). Consequently, the socio-economical effects may vary significantly between regions.

King scallops are known to exhibit high retention levels and low depuration rates (Blanco et al., 2002) contrary to most other species (Wohlgeschaffen et al., 1992; Álvarez et al., 2020), which can result in prolonged fishery closures. In the event of contamination, the ability to predict depuration dynamics would enable professionals to anticipate the potential contamination of king scallops with domoic acid at the start of the fishery season (from October to May in France) and to avoid costs that may outweigh the potential benefits. Knowing the level of contamination of shellfish before the opening date of fishing will allow fishermen to change their strategy in shifting to another species, postponing the beginning of the fishing season and the subsequent preparation of the vessel at a later date, or the modification of the fishing areas, depending on the capacities of the respective fleets impacted. In addition to their operational nature, the REPHYTOX and REPHY monitorings provide long-term data that allow to address the question of the inter-annual and spatial variability of domoic acid depuration rate in *P. maximus*. Through this original approach, we investigated whether depuration dynamics are mainly driven by environmental conditions or physiology. We also proposed an operational application of this knowledge by investigating how depuration, and thus domoic acid concentrations at the opening of the fishery season (autumn) might be predicted from contamination data acquired earlier in the year. To address these objectives, we identified and described *P. maximus* contamination events in each French province and then estimated the depuration rates for each contamination event. Based on these estimates, we assessed whether the depuration rates vary across provinces, time periods, maximum domoic acid concentrations and/or environmental conditions, or whether they appear independent from these factors. Then, we examined the potential to estimate the duration of a contamination event based on the initial domoic acid concentration and the average depuration rate. Finally, we studied the use of contamination data from other bivalves as sentinel species for the early detection of contamination.

## 2. Materials and methods

### 2.1. Studied provinces

For this study, we defined 13 provinces along the French Atlantic and the English Channel coastlines (Figure 1) based on knowledge on domoic acid issues and hydrodynamics; coordinates are provided in Supp. Table. S1. Domoic acid contamination events with sufficient data to investigate king scallop contamination occurred in 8 out of 12 provinces (names in bold in Figure 1); thus, only these 8 provinces will be presented in the Results section.

**Figure 1:**
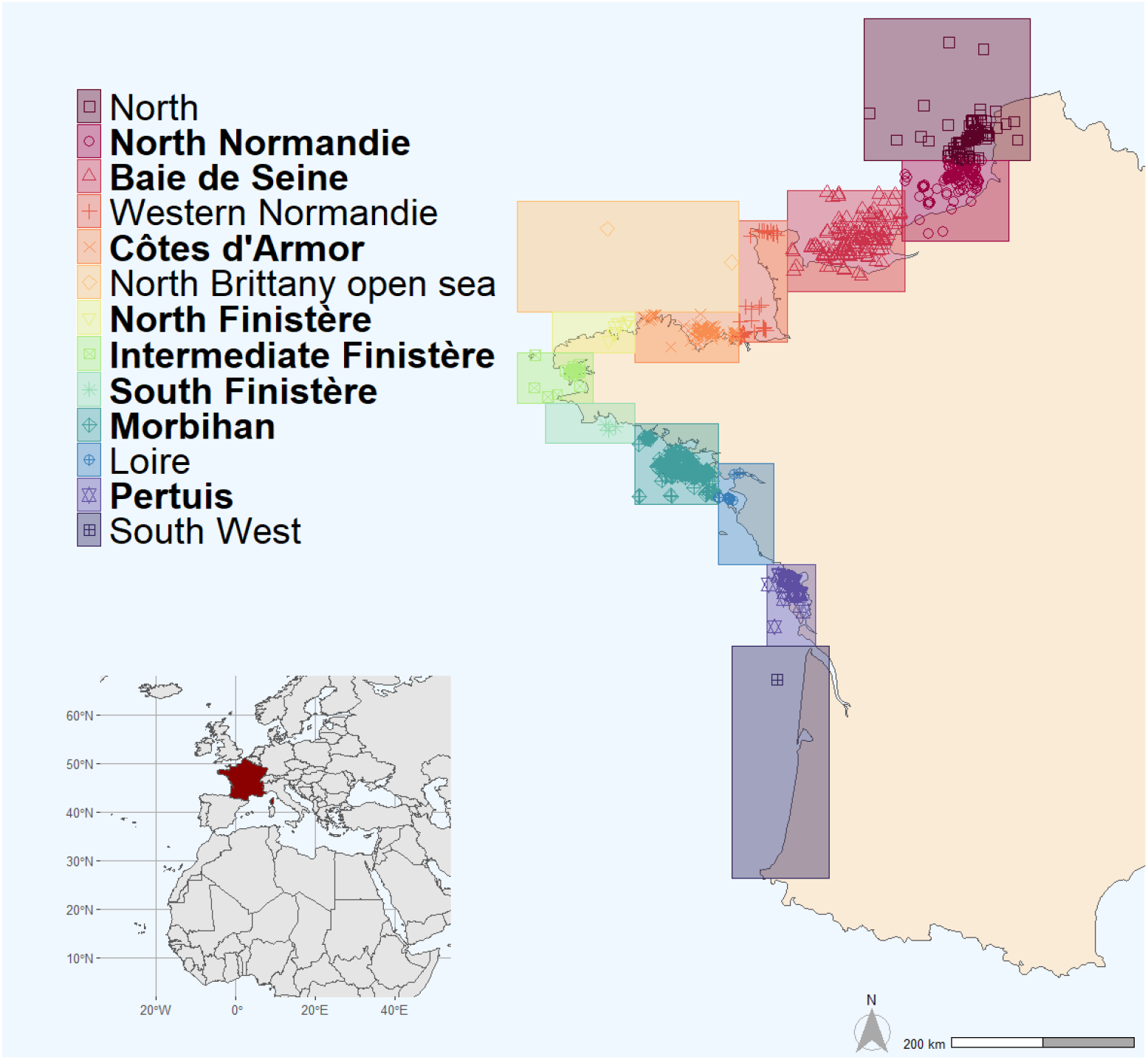
Map of the provinces studied for domoic acid concentrations in P. maximus and other shellfish species. The 8 provinces with sufficient domoic acid concentration data to study the depuration of *P. maximus* are in bold. Each dot represents one sampling location with available domoic acid concentrations recorded in the REPHYTOX dataset. The maps were edited using R, with the rnaturalearth package for the data and the ggplot2 and patchwork packages for the representation. Both maps are represented with the datum World Geodesic System 1984, and the world map is centered at latitude 52 and longitude 10, with France highlighted in red.

### 2.2. Data

#### 2.2.1. Data of domoic acid contamination in molluscs

Domoic acid concentrations in shellfish species were obtained from the Ifremer REPHYTOX programme initiated in 1987. Domoic acid measurements started in 1999 and are performed using HPLC-UV method, and quantification is expressed in mg of toxin per kg wet weight of total flesh. Detection and quantification limits range from 0.15 and 1.9 *mg kg*^−1^ depending on the analytical laboratory. The dataset of domoic acid concentrations from 1999 to 2024 was retrieved from the Quadrige database (https://envlit.ifre-mer.fr/Quadrige-la-base-de-donnees) in June 2024. The frequency of domoic acid measurement in shellfish varies by region and type of shellfish. For shell-fish offshore (*e.g*., king scallops, clams, common European bittersweet), monitoring generally involves samplings every fortnight during the exploitation period when toxin concentration is below the regulatory threshold. Other shellfish on the foreshore are only sampled when a proliferation of toxic microalgae is detected by the REPHY. The sampling is conducted every week if *Pseudo-nitzschia* spp. in the water column or toxin in shellfish are above the threshold. In addition to regular sampling for public health purposes, self-checks may be performed by fishermen, especially for *P. maximus* in the Bay of Brest. The dataset used in this study includes domoic acid concentrations for 17 molluscan shellfish species, not evenly represented. For instance, there are 6,858 data for *P. maximus*, 1,345 records for mussels *Mytilus* spp., and 664 for the common European bittersweet *Glycymeris glycymeris*. The other species have fewer occurrences of domoic acid measurement, as detailed in Table 1. The third column shows the number of occurrences with values exceeding the limits of detection and quantification. All analyses were conducted on R version 4.2.2 (R core team, 2022), and statistical metrics were computed per province and year of contamination (mean, standard deviation, standard error, minimal and maximal values).

**Table 1:**
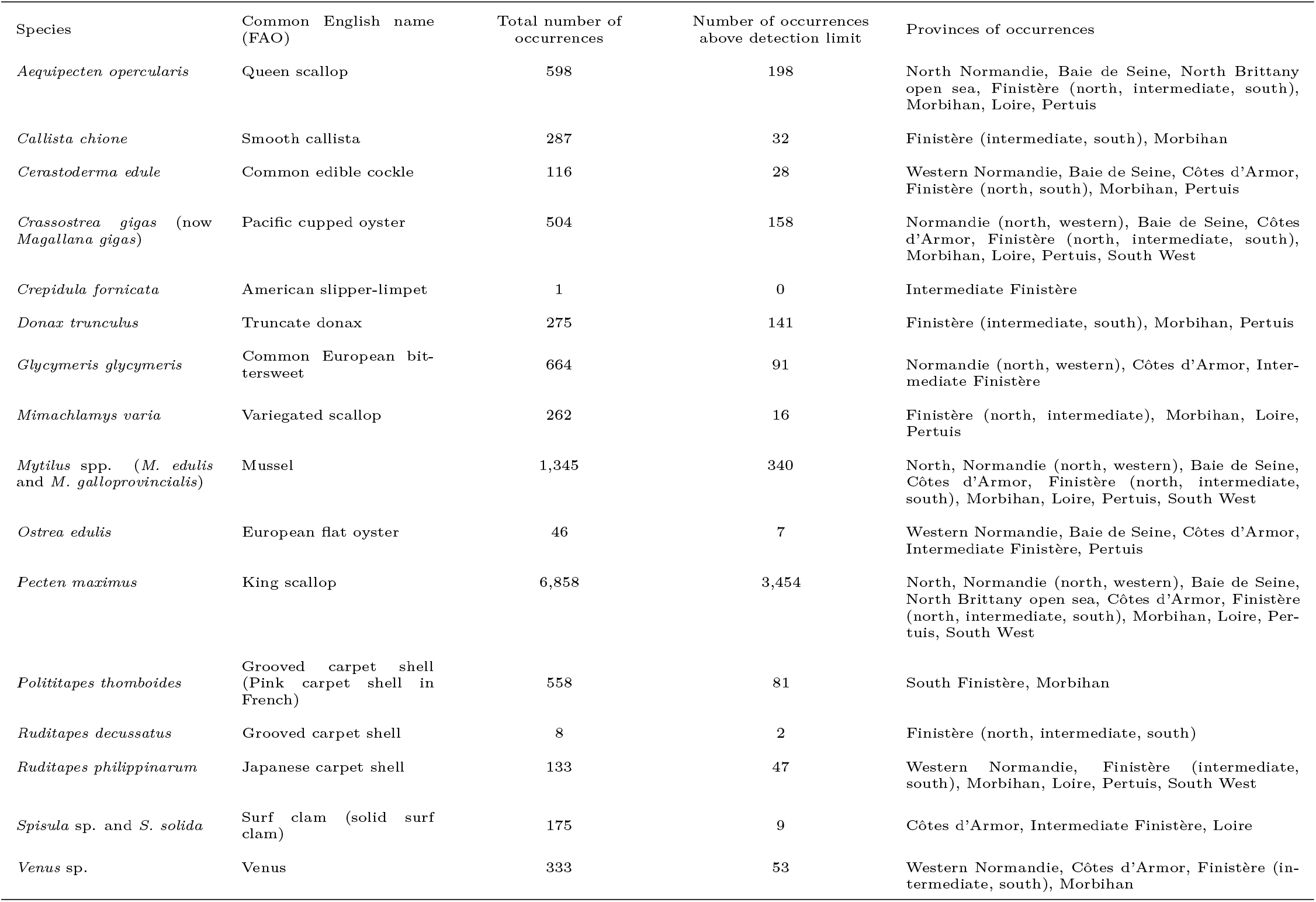
Number of domoic acid occurrences in tissues per shellfish species (given in alphabetical order) in the dataset considered and provinces where they are located. The English common names are given based on the FAO reference (https://www.fao.org/fishery/en/species).

#### 2.2.2. Environmental data

Environmental data were retrieved from the Ifremer REPHY programme initiated in 1987. The dataset was retrieved from SEANOE (www.sea-noe.org/) in which data are available in open-access (REPHY - French Observation and Monitoring program for phytoplankton and hydrology in coastal waters, 2023) and cover the period from 1987 to 2022. We considered 3 variables to describe the environmental conditions of the provinces, here given with their unit and code in the REPHY dataset: temperature (°*C*, “TEMP”), salinity (−, “SALI”) and chlorophyll-a concentration (*g L*^−1^, “CHLOROA”). For each province, a set of locations was selected and used to calculate the average environmental conditions: Normandie nord (At so, Dieppe and Dieppe 1 mile), Baie de Seine (Antifer ponton pétrolier, Cabourg, Ouistreham 1 mile, Roches de Grandcamp and St Aubin les Essarts); Côtes d’Armor (Dahouët, Trébeurden, Bréhat and Tréguier pont), Intermediate Finistère (Kervel, Kervel large and Lanvéoc), North Finistère (St Pol large), South Finistère (Concarneau large), Morbihan (Creizic, Lorient 16 and Men er Roue), Loire (Bois de la Chaise large and Boyard), and Pertuis (Nord saumonards).

### 2.3. Approach

#### 2.3.1. Contamination events in king scallops

For each location, domoic acid contamination events were identified only in *P. maximus* from 2004 to 2023. A contamination event was defined as a domoic acid concentration exceeding 20 *mg kg*^−1^ of total flesh weight (*i.e*., the European regulatory limit), followed by a decrease in concentration which corresponds to the depuration phase. The initial date, end date, and maximum domoic acid concentration for each event were reported. Thus, one contamination event can contain more than one occurrence as defined in Table 1 and 104 contamination events were described. The resulting dataset is available on SEANOE (Le Moan et al., 2025) and the scripts are available on https://versio.iuem.eu/eline.lemoan/domoic-acid-depuration-of-king-scallops.git.

#### 2.3.2. Depuration rate estimation, distribution and spatio-temporal variability

For each contamination event, the depuration rate was estimated by fitting an exponential decay model with a non-linear least squares regression performed in the R software (R core team, 2022) defined as: [*DA*]_*t*_ = [*DA*]_0_ × exp(−*τ* × (*t* − *t*_0_)), with [*DA*]_0_ the domoic acid concentration at initial time, *τ* the depuration rate, *t* the time at which we calculate domoic acid concentration and *t*_0_ the initial time. For further correlation analysis, only the fits corresponding to the following criteria were retained and named as “selected” events: *R*^2^ above 0.40 and more than 10 points distributed over the period (fits based on dots clustered at the beginning or the end of the time period were excluded). To test for differences between provinces and initial years, a one-way ANOVA was performed after checking normality and homoscedasticity of the residuals, followed by pairwise t-test using R (R core team, 2022). In cases of non-normality, the Kruskal-Wallis test was achieved, followed by Dunn’s test.

#### 2.3.3. Correlation between depuration rates and environmental conditions

First, correlation between maximum initial domoic acid concentration and depuration rates was tested. Then, to test for correlations between depuration rates and environmental conditions, such as water temperature, salinity, and chlorophyll-a concentration, we calculated the mean of each environmental variable over seven periods, based on the initial date of the depuration rate estimation. The seven periods are defined as follows:

- “Two-weeks”: between the initial date and the initial date plus 15 days;
- “Two-months”: between the initial date and the initial date plus 60 days;
- “Six-months”: between the initial date and the initial date plus 180 days;
- “Whole period”: between the initial date and the final date of contamination/decontamination rate estimation;
- “Spring whole”: between the 20^*th*^ of March and the 20^*th*^ of June of the year of the initial date;
- “Summer whole”: between the 21^*st*^ of June and the 23^*rd*^ of September of the year of the initial date, and
- “Autumn whole”: between 24^*th*^ of September and the 21^*st*^ of December of the year of the initial date.

For each relationship, a linear regression was performed between the depuration rates and the average per period of the environmental variables (temperature, salinity and chlorophyll-a concentration), using R software (R core team, 2022). The significance of the relationship was assessed by the p-value and the relevance of the relationship was determined by the adjusted *R*^2^. To determine correlations between variables, Spearman’s rank correlation test was used.

#### 2.3.4. Predictions of domoic acid depuration dynamics

The objective of this section was to assess the feasibility of predicting domoic acid depuration dynamics using only the initial domoic acid concentration, with the aim of determining domoic acid concentrations at any given time. Different scenarios were tested to improve the accuracy of the prediction: 1) using only the initial domoic acid concentration with the median depuration rate; 2) refining the date of the initial domoic acid concentration based on the end of the *Pseudo-nitzschia* blooms or taking into account recontamination; and 3) considering an inter-individual variability of domoic acid concentrations based on individual observations (García-Corona et al., 2024). The scenarios are detailed below, and the conditions of each scenario were added to the previous one. A prediction was defined as accurate if at least 50% of the measured domoic acid contamination points fell within this interval.

##### Application of the median depuration rate

First, we applied the median rate (0.0046 *d*^−1^) and the quantiles at 25% (lower limit, 0.0033 *d*^−1^) and 75% (upper limit, 0.0067 *d*^−1^), starting at the same initial value for the three depuration rates. This method was applied to the 104 events, and the number of points within this range was counted.

##### Refinement of the initial concentration value

Specifically, we tested scenarios based on the definition of the initial domoic acid concentration value used in the prediction. This refinement was based on two facts: (i) depuration kinetics are more accurate when considering contamination data at the end of the toxic algal bloom, when contamination is over, and (ii) potential recontamination by the presence of toxic algal cells is possible during the depuration period. Thus, in order to avoid erroneous estimation of the depuration rate, it is more accurate to divide the previously defined contamination events into several contamination events when recontamination occurs.

##### Considering inter-individual variability of domoic acid concentrations

There is high inter-individual variability in domoic acid concentrations, and data from the REPHYTOX programme correspond to the average of 10 individuals analysed as a pool. Thus, we made scenarios with a 10% interval around the initial domoic acid concentration to test the variability that could be observed in-situ. To test the robustness of our model, we choose the smallest variation that has been observed in previous results of king scallop domoic acid concentrations (García-Corona et al., 2024). We applied the median rate to the initial mean value that was adjusted in the previous step, the lowest rate to the highest domoic acid concentration (plus 10%) and the fastest rate to the lowest concentration (minus 10%).

##### Season of the initial concentration value

We assessed the optimal time of the year to sample king scallop tissue for domoic acid quantification for professionals, in order to evaluate the depuration kinetics and thus the potential closure or opening of the fishery activity at the beginning of the season in autumn. To this end, the events were divided into five categories depending on the month of their initial date: (i) January to February, (ii) March to May, (iii) June to August, (iv) September to November, and (v) December. The number of events accurately predicted for each category was counted. This procedure was repeated for the first scenario, using the median rate, and for the second scenario, with refinement of initial domoic acid concentrations.

##### Events unsuccessfully predicted

Under the previous scenarios, certain events proved challenging to be predicted, thus we identified specific cases for which the discrepancies between observations and predictions were relatively high. This approach was conducted in order to determine the conditions that presented the highest risk of inaccuracy in predictions, and consequently errors in the predicted date for opening of the fishery activity. These events include those with high depuration rate estimated, those with high variability in the data and those with variability in the initial domoic acid concentration.

#### 2.3.5. Predictions of dates when concentrations reach the regulatory threshold

As a second method, we focused on predicting when domoic acid concentrations would fall below the regulatory threshold of 20 *mg kg*^−1^, a key operational objective. To evaluate the predictive capacity of the approach, for each of the refined events (117 in total), as explained in section 2.3.4, a date was selected when the domoic acid concentration was close to the threshold and the most optimistic scenario was tested to compare the measured domoic acid concentration with the predicted one. This scenario assumed a 25% reduction of the initial domoic acid concentration (based on observed *in-situ* variability, see below section 3.2.1, Figure 3) and applied the fastest depuration rate from the upper quantile of the distribution (0.0067 *d*^−1^, see below section 3.2.1, Figure 3). The timing of the predicted fishery reopening has different consequences for professionals: on one hand, a delayed prediction may result in fishermen missing the fishing season due to a lack of boat preparation, whereas on the other hand, an early prediction could lead to unnecessary preparation of their boats without the ability to fish. This latter issue being less impacting for fishermen, we chose the optimistic scenario.

### 2.4. Sentinel species

The utilisation of other species present within the same geographical area as *P. maximus* to evaluate domoic acid contamination has the potential to facilitate the early detection of such contaminations, thereby enabling the anticipation of management strategies for fisheries. To assess the correlation between domoic acid concentrations in other shellfish and *P. maximus*, their concentrations were analysed in parallel within the same locations. For this study, and for shellfish species other than king scallops, we defined the “spring contamination” period as spanning from March 1^*st*^ to June 1^*st*^. For each province and each year, we recorded the number of data points during this period where other shellfish were either contaminated (domoic acid (DA) concentration above the regulatory threshold: [*DA*] ≥ 20 *mg kg*^−1^) or had detectable domoic acid levels ([*DA*] ≥ 1 *mg kg*^−1^). From these recorded events, we then counted the number of times during the fishing season (October 1^*st*^ to December 31^*st*^) where king scallops were either contaminated ([*DA*] ≥ 20 *mg kg*^−1^) or had detectable levels of domoic acid ([*DA*] ≥ 1 *mg kg*^−1^). Finally, the probability of detecting domoic acid or contamination in king scallops during the fishing season was calculated, taking into account the detection or presence above the regulatory threshold of domoic acid in other shellfish during the spring period.

## 3. Results

### 3.1. Contamination events of king scallops

Thanks to the analyses of domoic acid concentrations from REPHYTOX monitoring across all provinces, based on 6858 data, we defined 104 contamination events of king scallop to domoic acid from 2004 to 2024; *i.e*., a peak above or equal to 20 *mg kg*^−1^ followed by a decrease in concentration. The maximum domoic acid concentrations of these events ranged from 20 to 860 *mg kg*^−1^ (114 *±* 142 *mg kg*^−1^, *mean ± SD*) (Table 2). The intermediate Finistère province (*i.e*., Bay of Douarnenez and Bay of Brest) has both the highest number of domoic acid contamination events and the highest mean initial domoic acid concentration, as shown in Figure 2 (dark red). Intermediate Finistère and Morbihan are the provinces with the highest number of contamination events, with 41 and 26 events, respectively (Table 2). These events appeared almost every year, with 12 out of 20 years with a contamination event from 2004 to 2023 in Intermediate Finistère, and every year from 2005 to 2015 for Morbihan (Figure 2). Domoic acid contamination events are reported from March to December with two main periods where contaminations are reported (associated to sampling periods): May/June and September/October.

**Table 2:** Number of domoic acid (DA) contamination events in king scallops, in total and per province, and statistics of maximal DA concentration (in *mg kg*^−1^) of the contamination events that occurred in the different provinces: mean, standard deviation (SD), minimum (min) and maximum (max). The results are for the data from the REPHYTOX monitoring programme retrieved for the period between 2004 and June 2024.

| Province | Number of<br>DA contami-<br>nation events | Mean of the<br>maximum DA<br>concentration | SD of the<br>maximum DA<br>concentration | Lowest value<br>of the<br>maximum DA<br>concentration | Highest value<br>of the<br>maximum DA<br>concentration |
| --- | --- | --- | --- | --- | --- |
| North<br>Normandie | 1 | 85.1 | - | - | - |
| Baie de Seine | 14 | 95.4 | 87.9 | 23.4 | 303.8 |
| Côtes<br>d'Armor | 1 | 24.4 | - | - | - |
| North<br>Finistère | 4 | 98.1 | 86.1 | 23.4 | 177.8 |
| Intermediate<br>Finistère | 41 | 135.1 | 183.6 | 21.4 | 861.4 |
| South<br>Finistère | 11 | 96.9 | 49.4 | 62.6 | 237.0 |
| Morbihan | 26 | 105.8 | 143.9 | 20.4 | 484.0 |
| Pertuis | 5 | 131.4 | 82.5 | 41.9 | 213.0 |
| **All**<br>**provinces** | **104** | **114.2** | **142.3** | **20.4** | **861.4** |

**Figure 2:**
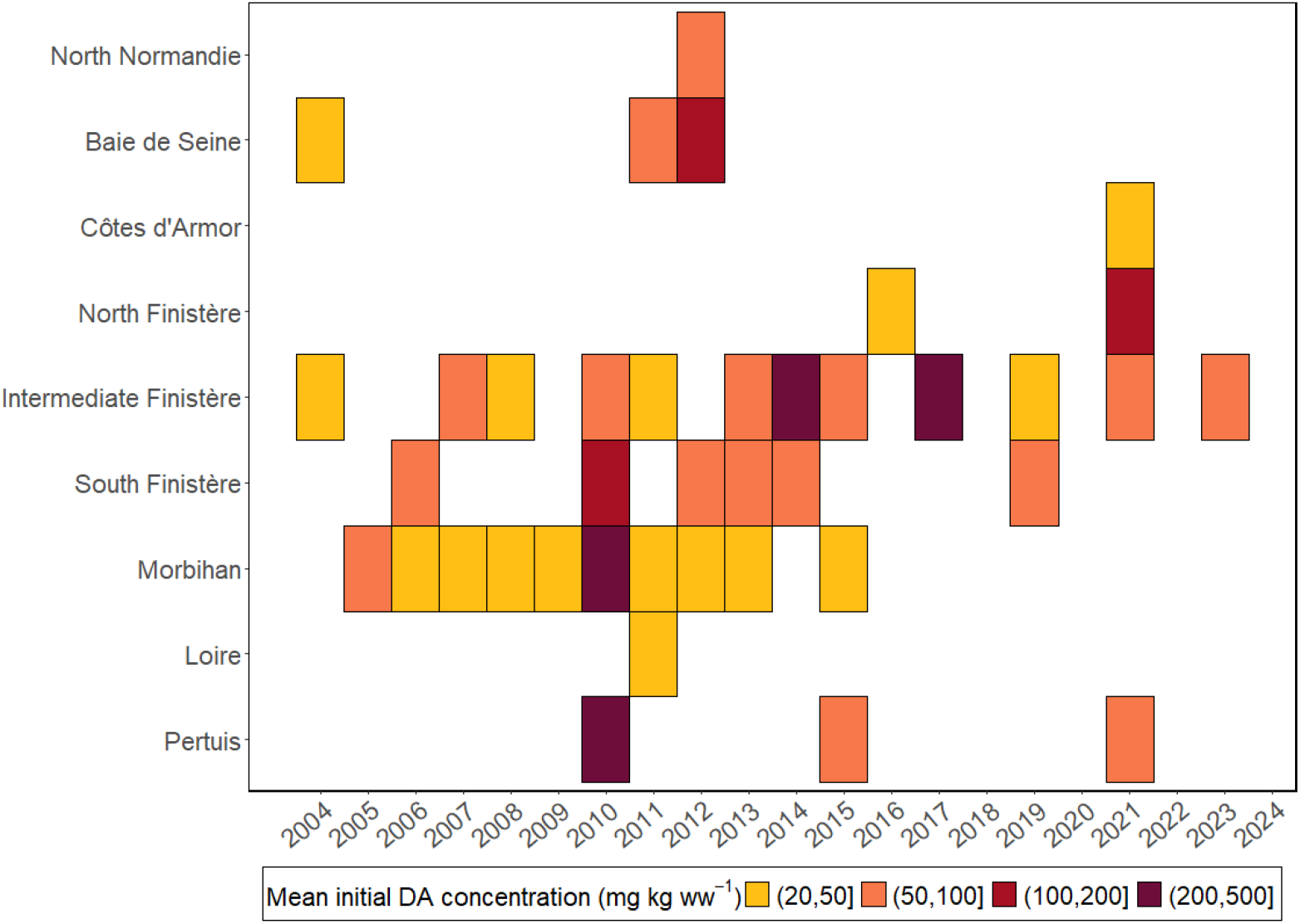
Heatmap with mean initial domoic acid (DA) concentration (*mg kg*^−1^) by year and by provinces in king scallops. The mean is calculated considering all the events in different locations in the same province for the same year. The colour scale shows the mean initial level of domoic acid concentration.

**Figure 3:**
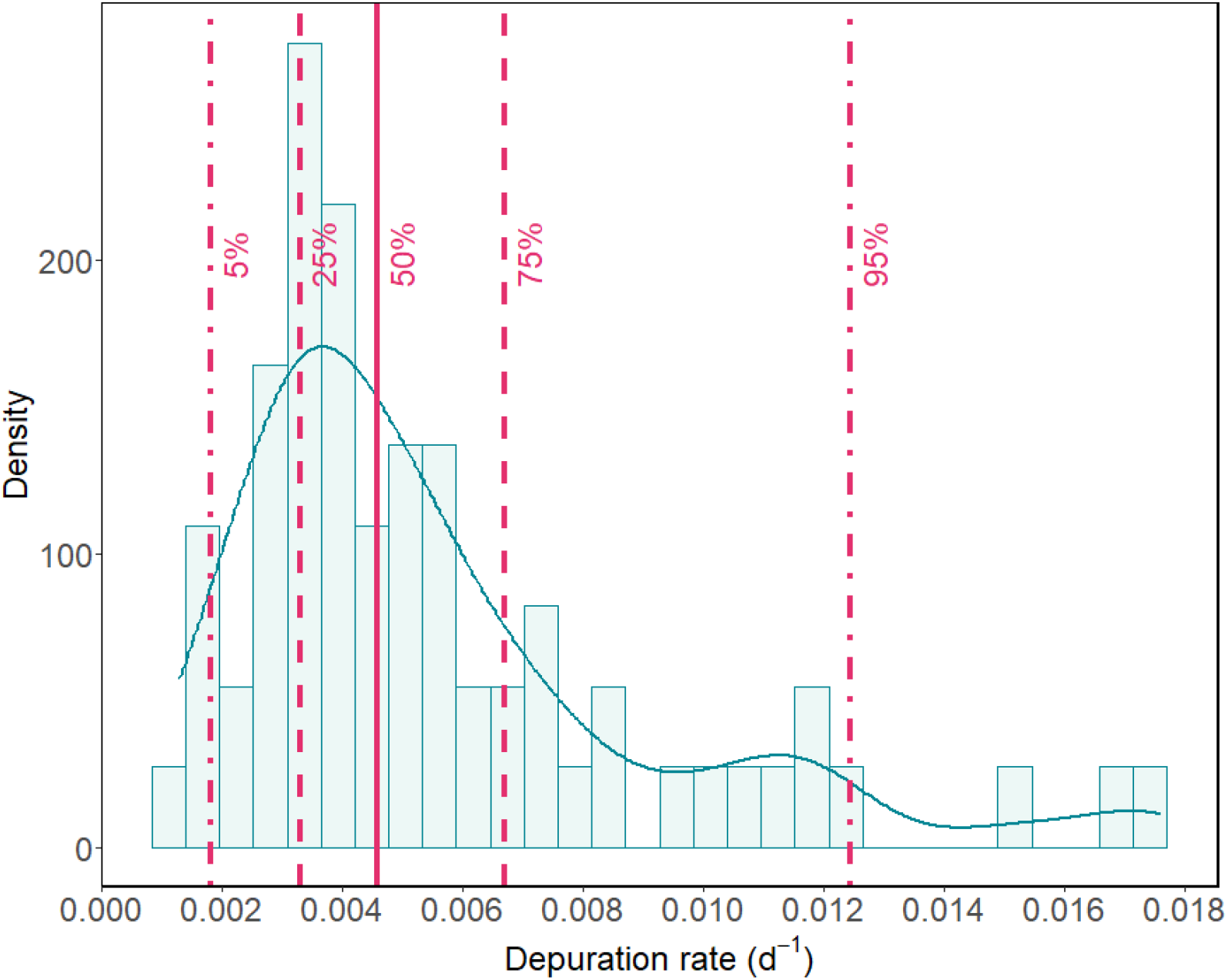
Density histogram of the depuration rates (*d*^−1^) with density function (blue curve) and quantiles (5%, 25%, 50%, 75% and 95%) as pink lines.

### 3.2. Depuration rate of king scallops

#### 3.2.1. Distribution of depuration rates

The depuration rates were estimated for the 104 events of domoic acid contamination in king scallops. After selection (section 2.3.2), we retained 63 depuration rates from 2004 to 2023 for further analysis. The distribution of these 63 selected depuration rates is presented in Figure 3 and shows that it does not follow a normal distribution, confirmed by Shapiro test (p-value *>* 0.05). However, the distribution is concentrated around a mean at 0.0057 *±* 0.0005 *d*^−1^ (*mean ± SE*) and the quantiles at 5, 25, 50, 75 and 95% were calculated giving depuration rates of 0.0018, 0.0033, 0.0046, 0.0067 and 0.0124 *d*^−1^ respectively.

#### 3.2.2. Variability of depuration rates

The variability of the depuration rates across years and provinces was assessed (Figure 4). Four outliers were identified: in 2005 at the Sud Belle-Ile station (Morbihan), in 2013 at the Golfe - la Teignouse station (Morbihan) and, in 2021 at the Gisement Roscanvel (Intermediate Finistère) and the Pertuis d’Antioche Coquilles Saint-Jacques (Pertuis) stations. The minimum estimated rate was found for the Gisement Moutons station (South Finistère) in 2006 and the maximum in 2021 for the Pertuis d’Antioche (Pertuis). The highest variability between the estimated depuration rates was observed for the Pertuis province with a standard deviation of 0.006 and a standard error of 0.003 for an average depuration rate of 0.009, higher than the global average (Table 3). The Baie de Seine and North Finistère are the provinces with an average depuration rate closer to the global average. The Kruskal-Wallis tests showed potential difference between provinces (p-value = 0.03) and no difference between years (p-value = 0.06). However, the Dunn pairwise tests showed no significant difference between provinces. The depuration rates for king scallops seemed to remain constant between periods and provinces from 2004 to 2023 with similar variability across provinces and years (Figure 4).

**Table 3:**
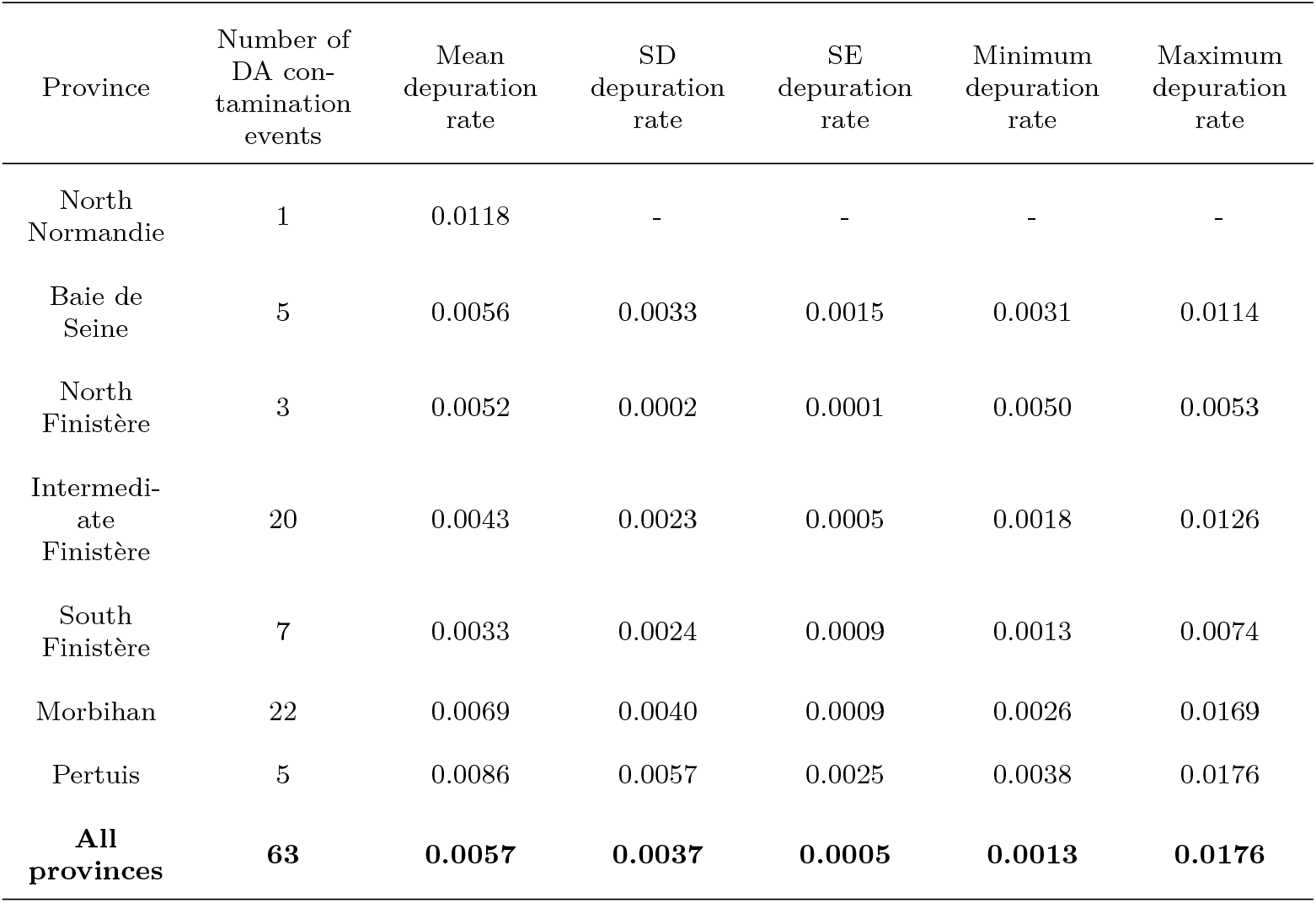
Summary of statistics of depuration rate (*d*^−1^) of the contamination events that occurred in the different provinces and all provinces grouped, with the number of domoic acid (DA) contamination events. Statistics are mean, standard deviation (SD), standard error (SE), minimum (min) and maximum (max) of the depuration rates.

| Province | Number of<br>DA con-<br>tamination<br>events | Mean<br>depuration<br>rate | SD<br>depuration<br>rate | SE<br>depuration<br>rate | Minimum<br>depuration<br>rate | Maximum<br>depuration<br>rate |
| --- | --- | --- | --- | --- | --- | --- |
| North<br>Normandie | 1 | 0.0118 | - | - | - | - |
| Baie de<br>Seine | 5 | 0.0056 | 0.0033 | 0.0015 | 0.0031 | 0.0114 |
| North<br>Finistère | 3 | 0.0052 | 0.0002 | 0.0001 | 0.0050 | 0.0053 |
| Intermedi-<br>ate<br>Finistère | 20 | 0.0043 | 0.0023 | 0.0005 | 0.0018 | 0.0126 |
| South<br>Finistère | 7 | 0.0033 | 0.0024 | 0.0009 | 0.0013 | 0.0074 |
| Morbihan | 22 | 0.0069 | 0.0040 | 0.0009 | 0.0026 | 0.0169 |
| Pertuis | 5 | 0.0086 | 0.0057 | 0.0025 | 0.0038 | 0.0176 |
| **All**<br>**provinces** | **63** | **0.0057** | **0.0037** | **0.0005** | **0.0013** | **0.0176** |

**Figure 4:**
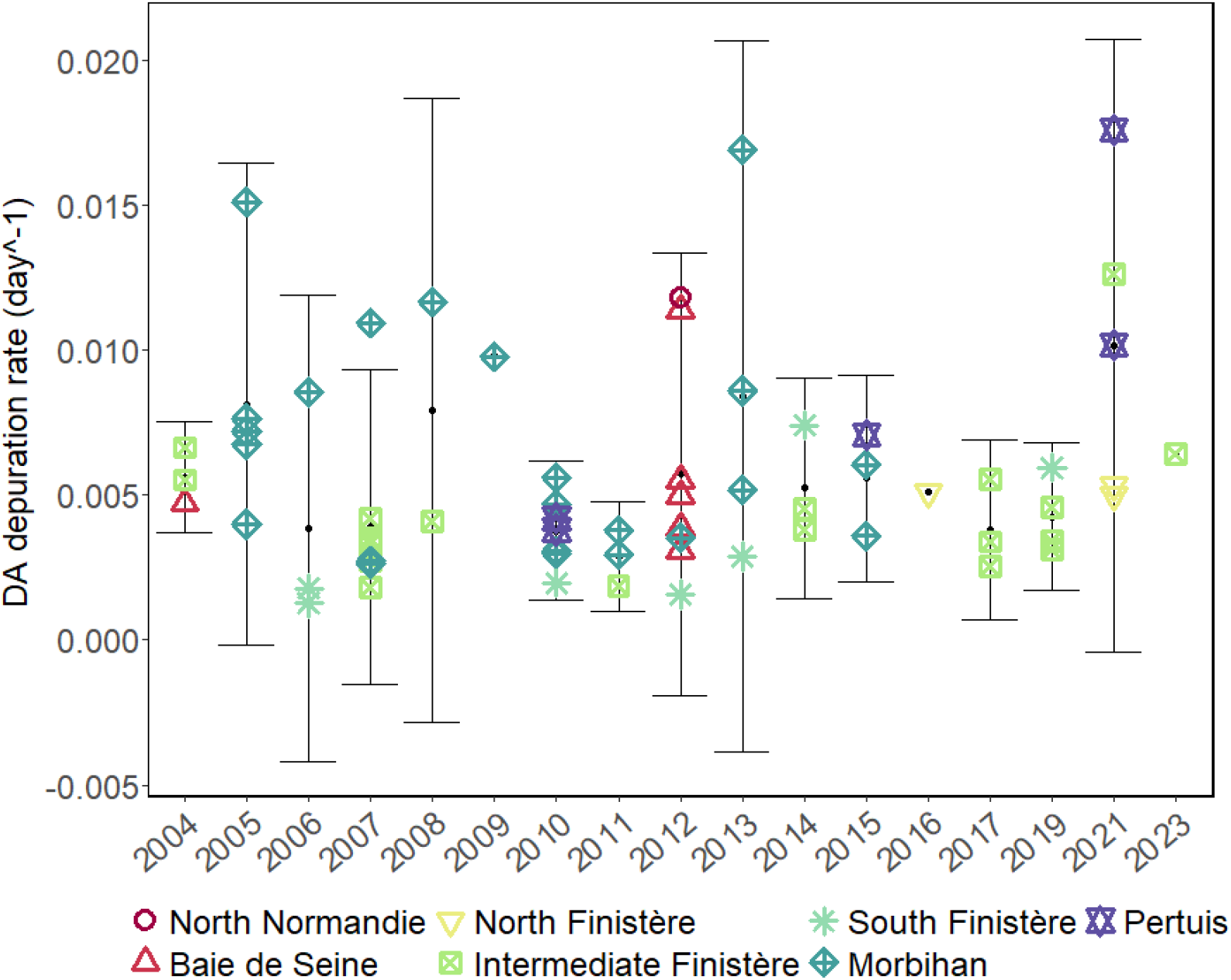
Domoic acid (DA) depuration rate of king scallops, *P. maximus* (*d*^−1^) over the years. The points (shape and colour) represent the provinces in which the depuration rates were calculated. Black dots represent the mean and the bars the standard deviation.

### 3.3. Correlation between depuration rate and environmental conditions

#### Maximum domoic acid concentration

The maximum domoic acid concentration recorded during each contamination event was compared to the corresponding depuration rate. No significant linear correlation was found between depuration rates and maximum initial domoic acid concentration (Figure 5), and no significant correlation was found with Spearman’s rank correlation test (p-value = 0.23), with a coefficient of 0.15 close to 0.

**Figure 5:**
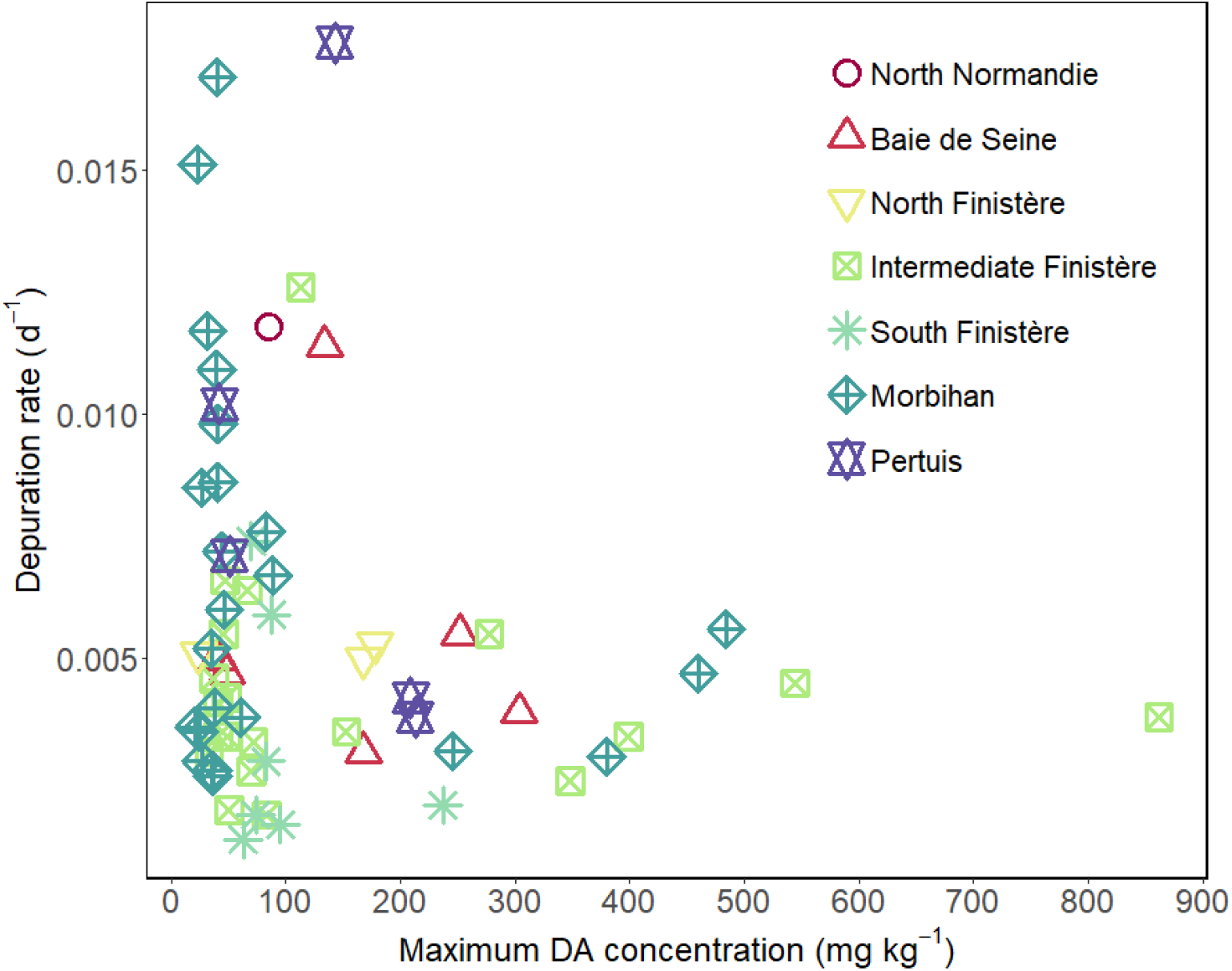
Estimated depuration rate (*d*^−1^) according to their corresponding maximum initial domoic acid (DA) concentration (*mg kg*^−1^) during a contamination event. This concentration was used as the initial value for the depuration rate estimation.

#### Environmental conditions

Correlations between depuration rates and environmental conditions were tested with linear models. The depuration rates were not correlated with environmental temperature when considering all locations. Indeed, based on the seven periods tested, only the summer period (from the 21^*st*^ of June to the 22^*nd*^ of September) showed significant correlation between temperature and depuration rates (p-value = 0.04), but it explained only 5% of the variance (Table 4). Depuration rates were correlated with salinity for the spring period (p-value = 0.001, from the 30^*th*^ of March to the 20^*th*^ of June), and for the summer period (p-value = 0.02), but it only explained 14% and 7% of the variance, respectively. No correlation was found between depuration rates and chlorophyll-*a* concentrations over the different periods considered.

**Table 4:**
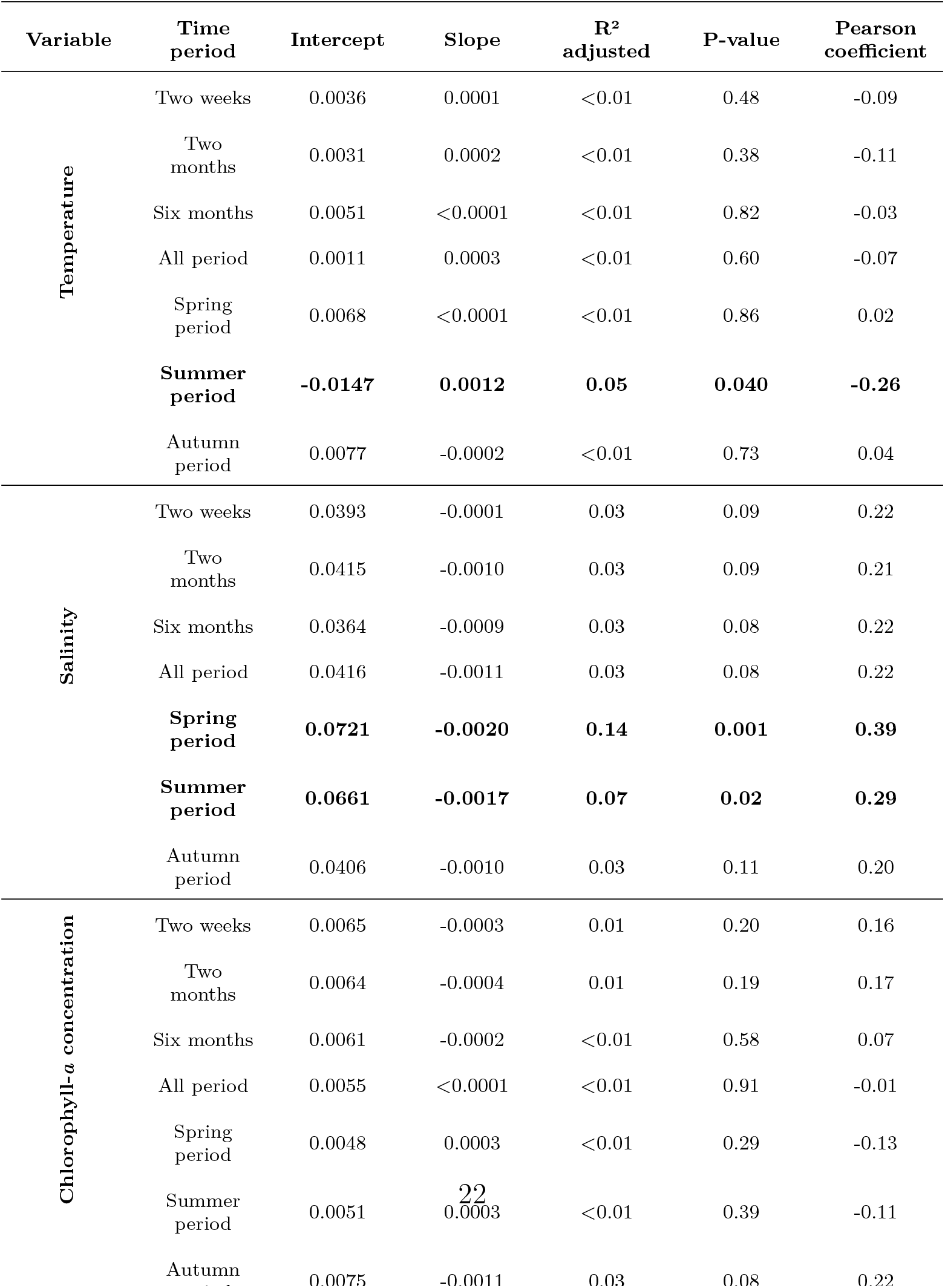
Coefficient of linear regression with p-value of the slope (Pearson correlation) and adjusted *R*^2^ for each environmental condition (sea surface temperature, salinity and chlorophyll-*a* concentration) and each period of time. The regressions were conducted on the selected depuration rates: the 63 values that satisfied the criterions of *R*^2^ above or equal to 0.4 and number of points above or equal to 10. The significant linear regressions are highlighted in bold in the table.

The depuration dynamic seems to be similar across all years, provinces and independent from environmental parameters, such as temperature, salinity and chlorophyll-*a* concentration. Thus, knowing the average domoic acid depuration rate for scallops, we can calculate an average depuration rate with confidence intervals to predict the depuration over time regardless of the environmental conditions.

### 3.4. Prediction of domoic acid depuration dynamics

#### 3.4.1. Application of the median depuration rate

Predictions of domoic acid concentration in king scallops over time were conducted on the 104 contamination events defined; including the 63 depuration rates “selected” (section 2.3.2) that were used to calculate the median rate, and the 41 “non-selected” events. In the aim of assessing if a single depuration rate could be used for operational predictions, and based on the distribution obtained in section 3.2.1, we computed predictions using the median rate and the rates at 25 and 75% of the distribution were used to determine the limits of the uncertainty range. In total, 47 events had 50% of their observations within the interval which represents 45% of satisfying predictions, an example is given in Figure 6A for the Pertuis Breton in the 2010 event.

**Figure 6:**
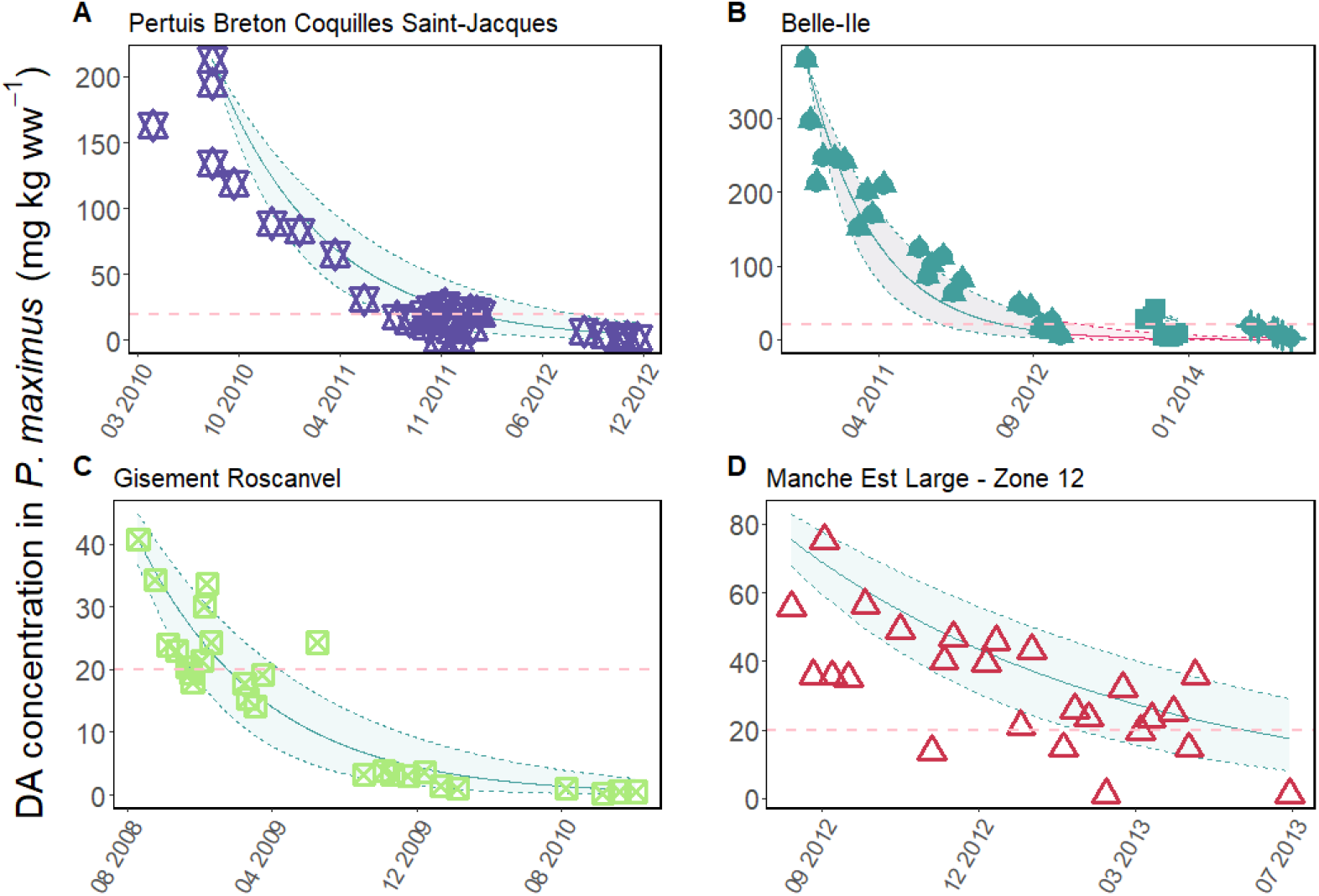
Observed and estimated domoic acid (DA) concentrations in king scallops, *P. maximus* according to time. Measurements come from the REPHYTOX datatset (coloured points according to the province) and estimations of the depuration (blue lines) with median rate (plain line) and higher and lower rates (dashed lines). Examples of depuration predictions in A) Pertuis Breton, a selected event for which the prediction gave more than 50% of points within the range, in B) Belle-Ile, an event for which the prediction was better when dividing the event to consider recontamination (blue line), the prediction without considering recontamination is represented in pink, in C) Gisement Roscanvel, a selected event for which the prediction was better with the 10%-interval around the initial value (associated with interindividual variability), and in D) Manche Est Large - Zone 12, a unselected event for which the prediction was better with the 10%-interval around the initial value

#### 3.4.2. Refinement of the initial concentration value

For 26 events, the initial domoic acid concentration used for the predictions was adjusted to consider depuration starting at the end of the contamination period and/or potential events of recontamination. This adjustment led to the consideration of 117 events instead of 104. After this adjustment, we obtained 56 events out of 104 (when considering only the end of the bloom), and 63 out of 117 events (considering both end of bloom and recontamination, thus leading to the division of several of the previous events), which provided more than 50% of their points within the range of depuration kinetics. This adjustment allowed for 54% of good predictions such as the example in Figure 6B for Belle-Ile from 2010 to 2015. Adjusting the initial concentration value thus increased the number of good predictions by 10%.

### 3.5. Considering inter-individual variability of domoic acid concentrations

To account for inter-individual variability in domoic acid concentrations and to increase our ability to represent depuration dynamics, we considered a variability around the initial domoic acid concentration of 10%. When considering this interval, we obtained a higher number of events with at least 50% of their points within the range of predictions, corresponding to 79 out of 117 events. Examples of satisfying predictions are shown in Figure 6C and D with selected and non selected events respectively, as explained in section 3.4.1. Thus 68% of events were well predicted. Considering inter-individual variability with 10% around the initial domoic acid concentration value thus increased the number of good predictions by 14%.

#### 3.5.1. Season of the initial concentration value

In order to better determine the best sampling period to quantify domoic acid in king scallop tissues for operational purposes, we evaluated the number of good predictions depending on the month at which the initial value was provided in the dataset for the decontamination event. We grouped them in 5 categories: the initial value for contamination occurred between January and February, in the spring from March to May, in the summer from June to August, in autumn from September to November and in December. Results showed that the predictions were better when the initial contamination value was given in spring (Table 5), whereas the lower number of good predictions occurred when the initial contamination value was recorded in autumn. The predictions based on summer initial points provided on average around 50% good predictions. When the initial contamination value is in December, the predictions are very good, however this corresponds to a small number of events; *i.e*. 6 events.

**Table 5:**
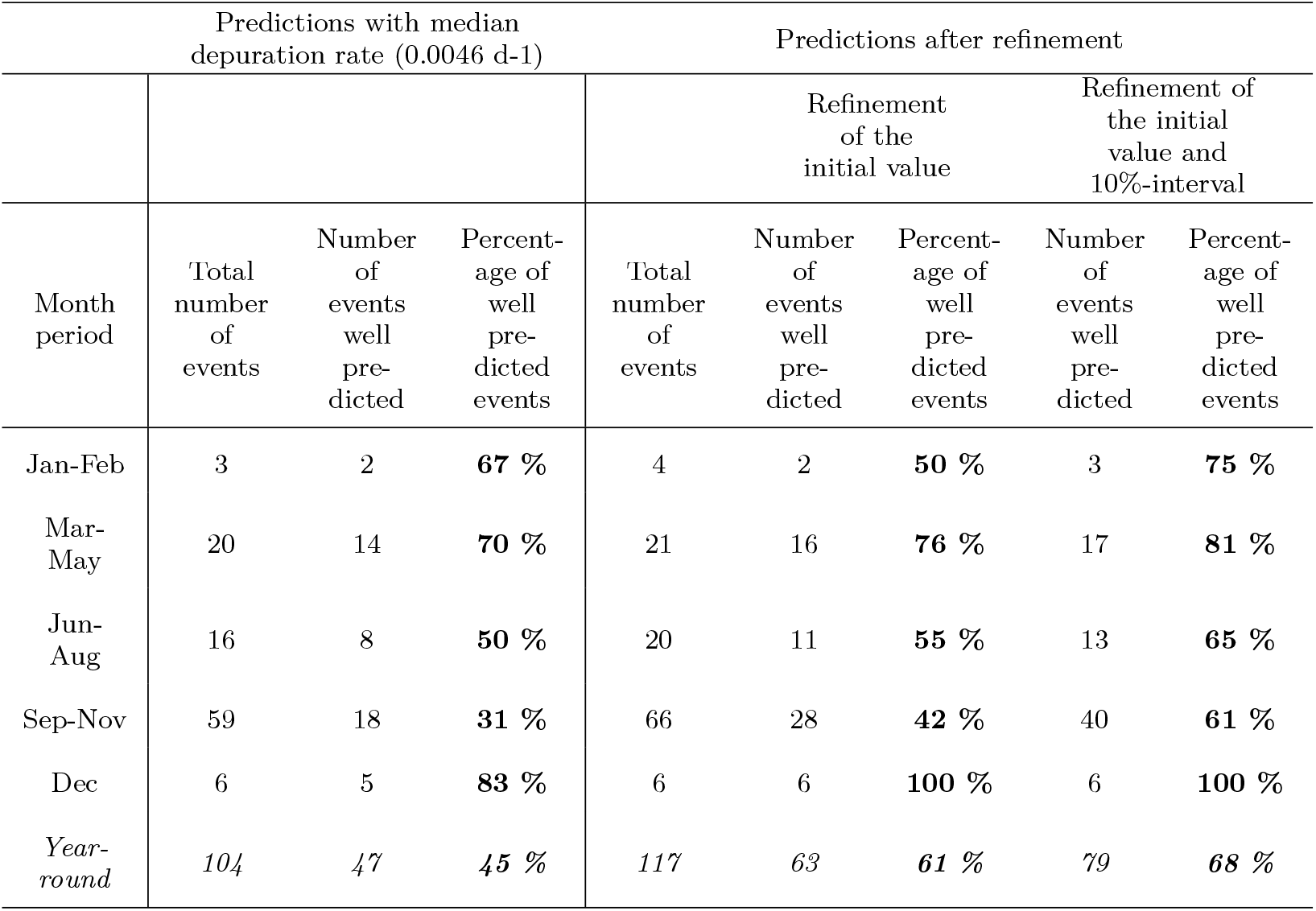
Number of domoic acid (DA) contamination in king scallop, *P. maximus* events classified by the month of their initial DA concentration value, and number of events with more than 50% of their points within the range of predictions considered as well-predicted events, and percentage of these well predicted events per category of month of the initial concentration value. These numbers and percentages are given for the three scenarios: application of the median depuration rate, refinement of the initial domoic acid concentration value and addition of a 10%-interval around the refined initial value to consider for interindividual variability.

### 3.6. Events unsuccessfully predicted

As shown in Table 5, several events were not successfully predicted, indicating a discrepancies between the estimated and observed depuration rates.

Thus, we defined the different specific cases for which predictions are difficult, to identify the limits of the method to be able to develop a reliable predicting tool.

Our first specific case is for events that showed a high depuration rate, which did not allow to make a good representation with the median and quartiles depuration rates. This is the case for one location in particular, Quiberon-concessions, for which the predictions were slower in 2006, 2007, 2008 and 2009 leading to a latter prediction of reopening of the fishery, while they were almost perfect for 2010 and 2015 (Figure 7A).

**Figure 7:**
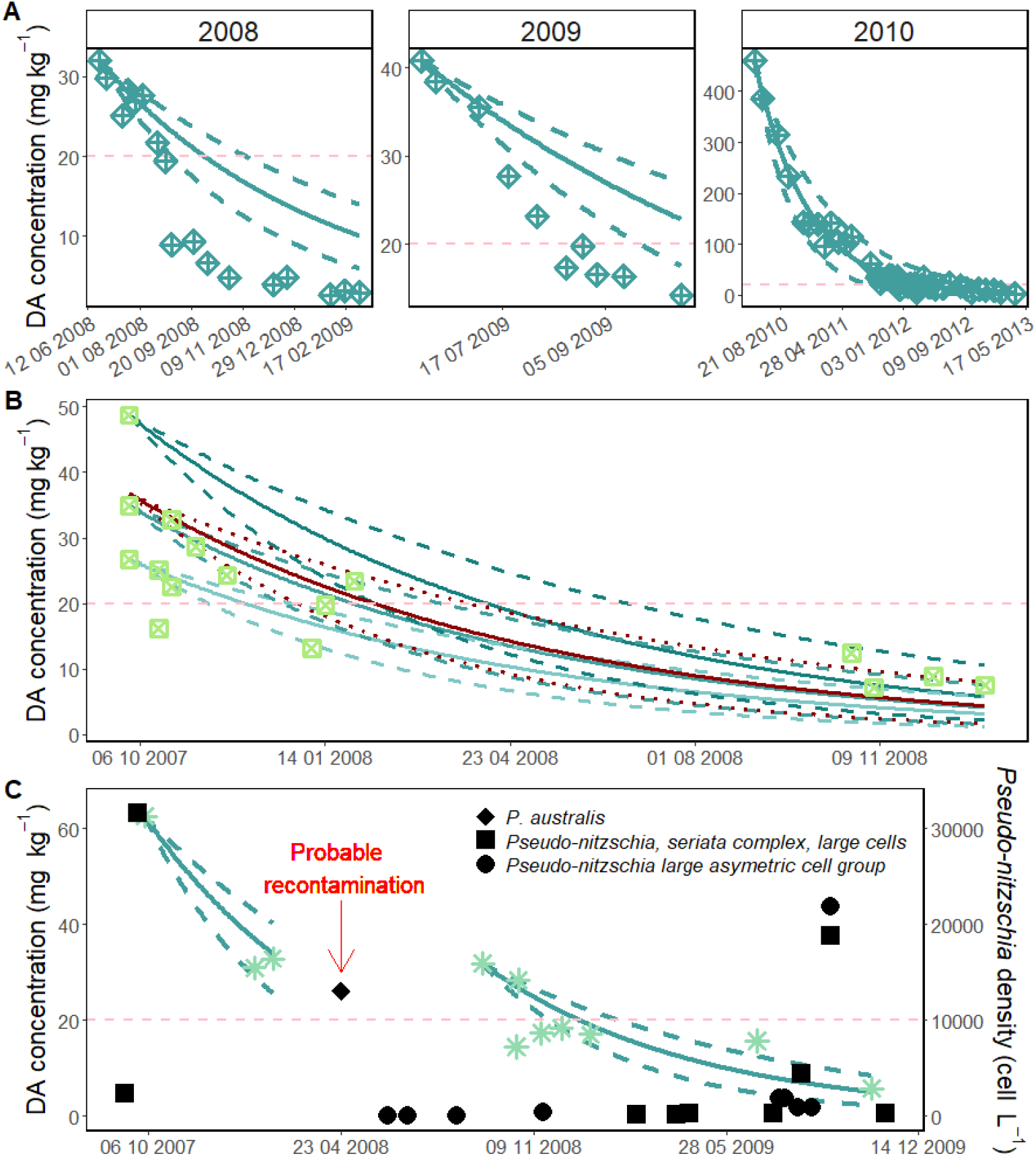
Domoic acid (DA) concentration (in *mg kg*^−1^) in king scallop, *P. maximus* tissues according to time with predictions of concentrations using exponential decay model to represent several examples of difficulties to predict the depuration dynamics. Predictions are defined with median depuration rate (plain blue lines) and with the depuration rates at 25% and 75% of the distribution (dashed blue lines) starting at the maximum initial domoic acid concentration. Specific examples for A) the site of Quiberon-concessions in three following years (2008-2010), B) the site of Gisement Ouessant in 2007 with red lines starting as the mean of the 3 different domoic acid initial concentrations and C) the site of Gisement Moutons in 2007, black dots correspond to the cell abundances of *Pseudo-nitzschia* species or groups recorded by the REPHY in the South Finistère province (Concarneau large, Amont port Kerdruc Rosbras and Estuaire amont Isle). Each taxonomic group is represented with a specific dot shape, including the most toxic known group being *P. australis* (the other groups comprised *P. subpacifica, P. fraudulenta* and *P. seriata*). The horizontal pink dashed line represents the regulatory threshold of 20 *mg kg*^−1^.

The second specific case was highlighted by a variability in the initial domoic acid concentration measured at the same or close date. The Gisement Ouessant in 2007 is an example (Figure 7B). For the initial date, 3 different values of concentration were recorded with 36.8 ± 11.1 *mg kg*^−1^ (*mean* ± *sd*) and for which we applied our median depuration rate, thus providing 3 different scenarios (Figure 7B, green lines). Finally, considering the mean value of the 3 concentrations provided a better prediction than considering each point separately as the initial value (Figure 7B, red line).

The last case that makes prediction difficult is the presence of potential recontaminations by the presence of *P. australis* in the environment. This can be demonstrated by the Gisement Moutons in 2007 (Figure 7C). For these events, considering one long-term event does not allow to accurately predict the observed depuration dynamic using the median depuration rate, it was necessary to separate each small recontamination event. To highlight the link between *P. australis* presence and domoic acid contamination in king scallops, density of *Pseudo-nitzschia* cells at the same period and for the same province using REPHY dataset was added to the depuration dynamic for Gisement Moutons in 2007 (Figure 7C).

### 3.7. Prediction of concentrations close to regulatory threshold

The second key operational objective was to assess our ability to predict domoic acid concentrations near the regulatory threshold under the most optimistic scenario, providing insight into the potential reopening of the fishing activity. The predictions were accurate for 61 contamination events (52%, Figure 8A), among which 23 contamination events (19.7%) presented low domoic acid values with initial value close to the 20 *mg kg*^−1^ regulatory threshold, which obviously led to accurate predictions (Figure 8B). Additionally, in 34 cases (29.1%), predicted domoic acid concentrations were lower than the observed values at the time they passed under the regulatory threshold (Figure 8C).

**Figure 8:**
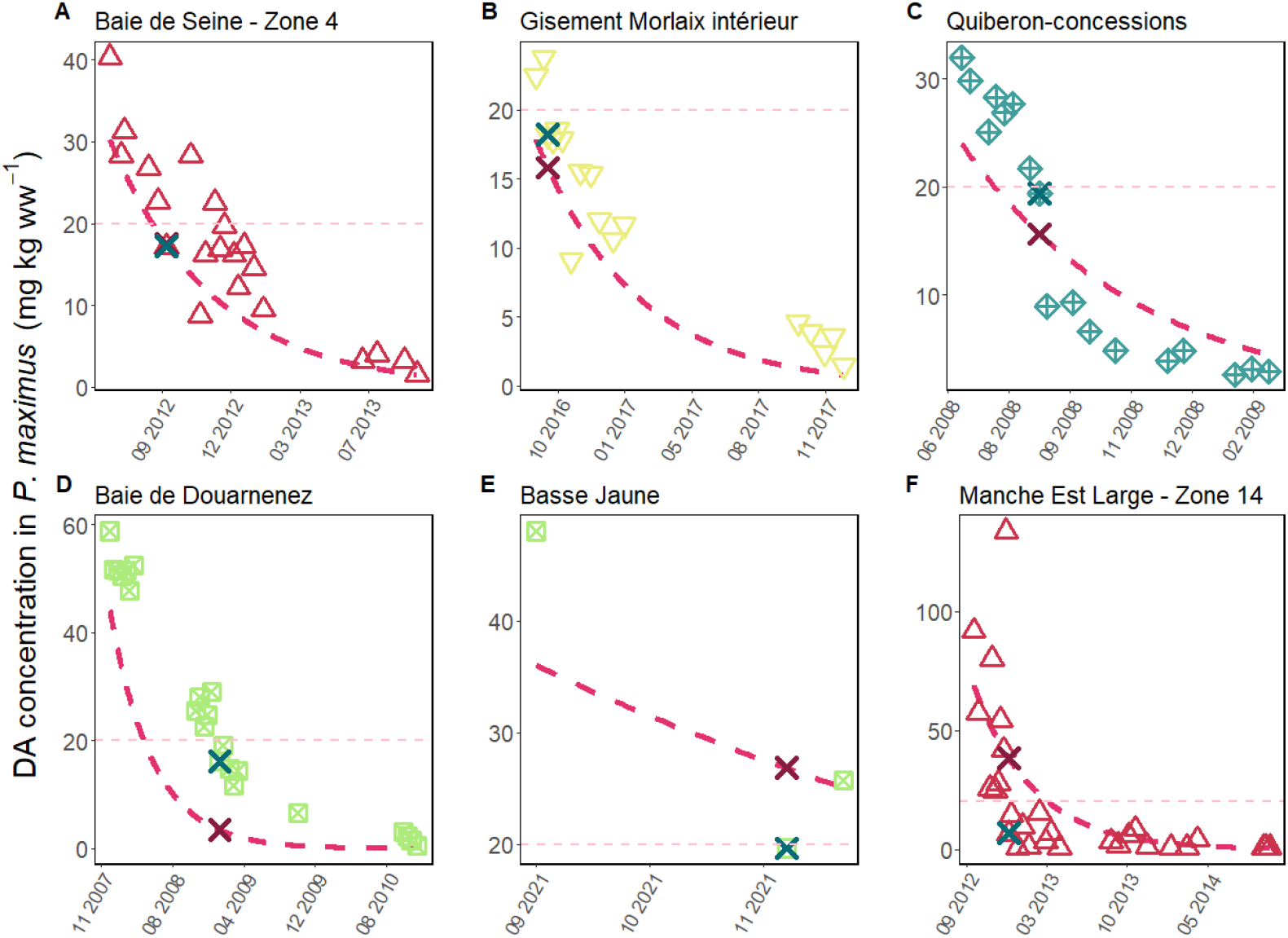
Domoic acid (DA) concentrations in king scallops, *P. maximus* according to time, recorded in the REPHYTOX dataset (points with shape and color according to the province) and predictions of depuration with the optimistic scenario (pink dashed line). The domoic acid concentration measured close to the 20 *mg kg*^−1^ threshold is represented with the blue cross, and the predicted one at the same date as the pink cross. The examples of predictions with the optimistic scenario are given for the six cases defined: A) good prediction, B) low concentration, C and D) opening predicted before the real date, with low (< 6 months) and high (> 8 months) difference, E) cases that are close but opening is predicted after the real date and F) cases that did not work.

However, 22 events (18.8%) were not well predicted. Among these, 7 events (6%) showed an overly rapid predicted depuration (Figure 8D), leading to a forecast reopening date much earlier than the actual observed one. In another 5 events (4.3%), the predictions were inaccurate but close, with the predicted concentration dropping below the 20 *mg kg*^−1^ threshold shortly after the observed one or with a low number of data points which do not go below the regulatory threshold (Figure 8E).

The last 10 events difficult to predict (example in Figure 8F) are detailed in Figure 9, including 5 events in the Bay of Seine (Figure 9A-E) which exhibits a high variability in the measured domoic acid concentrations. In this province, the geographical zones are large and the samples are not taken at the same place according to GPS coordinates in the REPHYTOX dataset. The variability in the data was also high for the two events in Morbihan (Figure 9F, G) and the 2021 event in Intermediate Finistère (Figure 9H). In a second event in Intermediate Finistère in 2023 (Figure 9I), the sampling was conducted by two different persons (fishermen) with the switch in mid-October 2023. In the Pertuis d’Antioche in 2021 (Figure 9J), the averaged temperatures recorded in summer and autumn periods (REPHY data) were the highest compared to all the other events of contamination.

**Figure 9:**
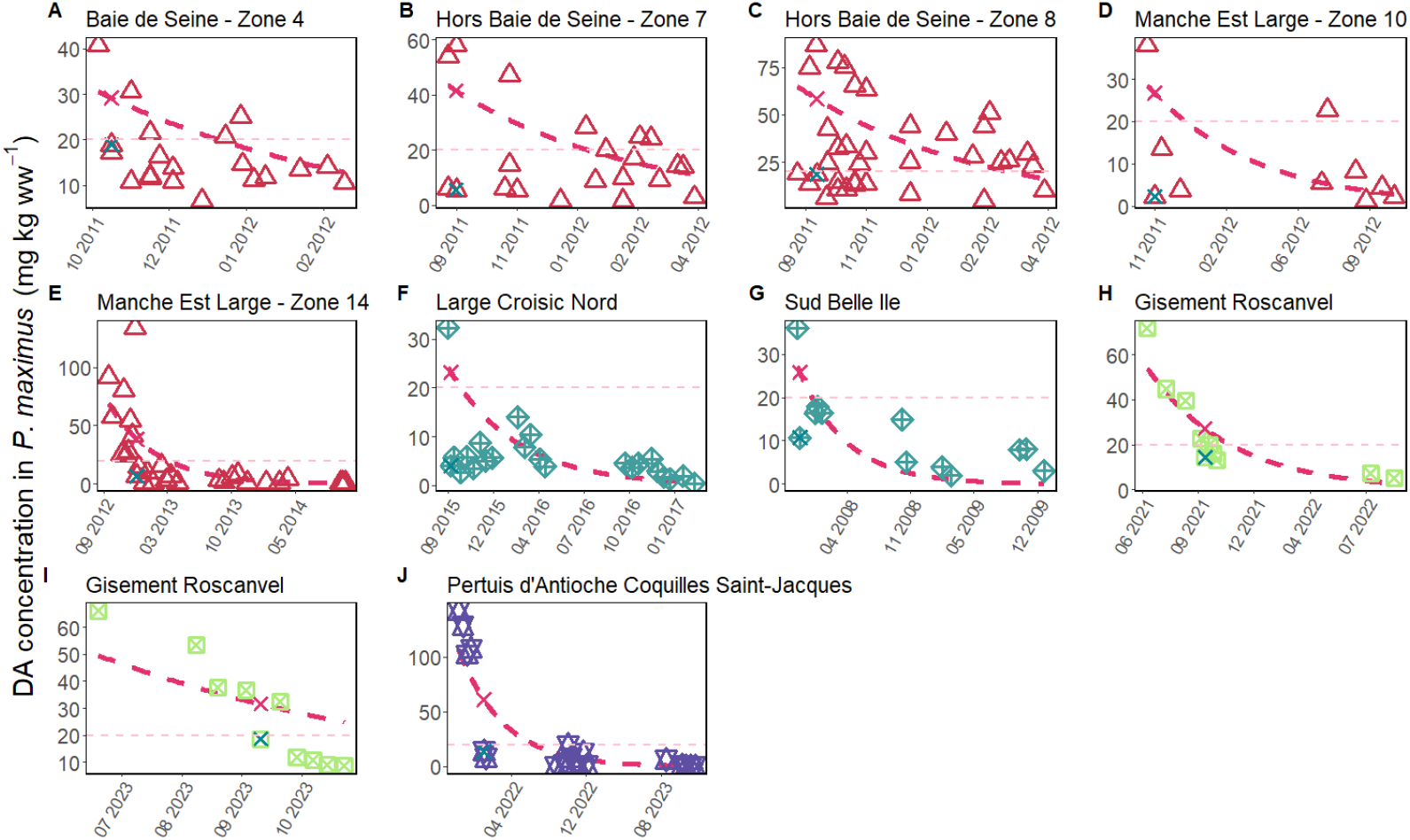
Domoic acid (DA) concentrations in king scallops, *P. maximus* according to time recorded in the REPHYTOX dataset (points with shape and color according to the province) and predictions of depuration with the optimistic scenario (pink dashed line). For the prediction of the concentration close to the regulatory threshold; the measured domoic acid concentration close to the 20 *mg kg*^−1^ threshold is represented with the blue cross, and the predicted one (at the same date) as the pink cross. The examples are the 10 specific cases for which predictions were not accurate, with from A to E, the Bay of Seine province, F-G the Morbihan, H-I the intermediate Finistère and J the Pertuis.

### 3.8. Sentinel species

Analysing side by side domoic acid concentrations of king scallops and other shellfish species in the same locations gave us insight into the use of other shellfish species as sentinel species. When looking at domoic acid detections (concentration above 1 *mg kg*^−1^) of other shellfish species during spring, 119 domoic acid detection in other shellfish corresponded to 107 detections (90%) of domoic acid in king scallops (≥ 1 *mg kg*^−1^), among which 42 contaminations (35%) above the regulatory threshold (≥ 20 *mg kg*^−1^), in the same locations at the time of the king scallop fishing opening season (October). When looking at contaminations of other shellfish (≥ 20 *mg kg*^−1^) during spring, 15 contaminations of other shellfish corresponded to 13 contaminations (85%) of king scallops, in the same locations, at the time of the king scallop fishing opening season (October).

## 4. Discussion

### Domoic acid contamination events for the last 20 years in 9 French provinces

This study investigated 20 years of data from the Atlantic and English Channel coastlines of France highlighting the value of long-term monitoring programmes not only for public health purposes but also for advancing our understanding of species-specific mechanisms of domoic acid contamination and depuration. A total of 13,422 records of domoic acid concentrations in shellfish tissues were considered, including 6,858 observations for *P. maximus*. From this dataset, we identified 104 contamination events in *P. maximus*, characterised by toxin concentration exceeding the regulatory threshold of 20 *mg kg*^−1^ in total tissues, followed by a subsequent decrease. These events provided a unique opportunity to investigate the depuration kinetics of domoic acid in *P. maximus* under natural conditions. To our knowledge, this is the first study to estimate depuration rates from such a large number of contamination events and to assess their relationships with environmental conditions. Furthermore, we successfully developed predictive statistical models for domoic acid concentrations in *P. maximus* based on initial concentration levels, offering potential operational applications for risk assessment and fishery management as discussed later.

The spatial distribution of contamination events revealed regional disparities in France, with certain areas experiencing recurrent domoic acid accumulation in king scallops, while others remaining largely unaffected. The Intermediate Finistère province (*i.e*. Bay of Douarnenez and Bay of Brest) accounted for 40% of the recorded contamination events (41 out of 104), with a maximum domoic acid concentration of 861 *mg kg*^−1^ in *P. maximus* tissues (recorded in April 2014 in Gisement L’Auberlac’h following a massive bloom of P. australis). The Morbihan province followed with 26 events and a maximum concentration of 484 *mg kg*^−1^. Together, these two provinces, located in Brittany, represented 64% of all recorded *P. maximus* domoic acid contamination events. Due to the high frequency of occurrences, these regions experienced domoic acid concentrations exceeding the regulatory threshold of 20 *mg kg*^−1^ over numerous years between 2004 and 2024. In contrast, surrounding regions, such as northern Brittany and the Pertuis provinces were only sporadically affected by contamination events. Understanding the drivers of *Pseudo-nitzschia* proliferations remains challenging and has been previously studied by (Husson et al., 2016) using similar datasets from the REPHYTOX and REPHY monitoring programmes between 1996 and 2012. Their study is based on *Pseudo-nitzschia* observations, whereas ours is based on domoic acid concentrations in *P. maximus* tissues. However, the spatial patterns of bloom frequency reported are consistent with our findings. Similar investigation of prevalence of domoic acid in bivalve tissues was conducted in Galicia by (Blanco et al., 2021) who investigated a 25-years dataset of domoic acid concentrations in several shellfish species. As expected, *P. maximus* was the species with the highest number of data and the highest domoic acid concentration. This species always contained domoic acid, thus it was not relevant to conduct any prevalence analysis on this species. Therefore, they focused on the second most contaminated species, the mussel *Mytilus galloprovincialis*, to be used as a sentinel species. They found spatial variations in terms of events and maximum domoic acid concentrations. Their results highlighted the geographical heterogeneity in domoic acid contaminations, emphasising the need for region-species management strategies.

As reviewed in (Lelong et al., 2012), in European waters, *Pseudo-nitzschia* sp. blooms mainly occur between January and May, as commonly observed in the Bay of Brest for example. This bloom period corresponds to our data which showed that contaminations of *P. maximus* mainly occur in spring. However, along the French Atlantic and English channel coastlines, Husson et al. (2016) found that *Pseudo-nitzschia* blooms occur from March to November with a peak from May to September, which is later than what we observed in our domoic acid contamination data. Nevertheless, since the REPHYTOX monitoring is based on fishing activities, data of domoic acid contamination in king scallops are biased by the methodology of sampling with regard to the time period and the frequency. For king scallops, the fishery season starts in October and finishes between March and May depending on the zone. The data are linked to the method used in the monitoring programme. For example, the French monitoring REPHYTOX is based on the sampling of commercial shellfish during the fishing season and additional sampling if there is the presence of potentially toxic phytoplankton species recorded by the REPHY. Other monitoring methods may be based on sentinel species which lead to sampling in other species in case of concentrations exceeding a defined threshold, or on specific demand from professionals, such as in Galicia, where fishing activity of king scallops is closed but sampling can be asked to assess if the concentration is suitable for fishing and selling during a specific and short period. Thus, all results linked to the season of the data need to be carefully interpreted according to this bias.

### Small variability in domoic acid depuration rates in *P. maximus*

After identifying the contamination events across the provinces, we investigated the depuration dynamics of *P. maximus*. The apparent domoic acid depuration rates were estimated using an exponential decay model, as commonly found in the literature to estimate domoic acid depuration rates (Blanco et al., 2002; Mafra et al., 2010; Álvarez et al., 2020) as well as for other phycotoxins such as paralytic shellfish toxins (Blanco et al., 1997). The average depuration rate estimated from 63 contamination events in *P. maximus* was 5.7 ± 10^−3^ 0.5 10^−3^ *d*^−1^ (*mean* ± *SE*), aligning with the findings of Blanco et al. (2002), who reported a depuration rate of 0.0078 *d*^−1^ using a similar exponential decay model on data obtained from experimental depuration monitored under controlled conditions. We also observed higher depuration rates in certain events, reaching up to 0.0176 *d*^−1^, comparable to the findings of Blanco et al. (2006), who found a maximum rate below 0.011 *d*^−1^ for whole flesh. However, even the highest rates observed in *P. maximus* remained lower than those reported for fast depurators, such as *Argopecten purpuratus* (0.27 *d*^−1^, under natural conditions, Álvarez et al. (2020)) or even *Placopecten magellanicus*, a relatively slow depurator (0.1 *d*^−1^, under experimental conditions, Douglas et al. (1997)). However, the depuration rates estimated in this study correspond to apparent rates because they may consider both uptake and elimination occurring concurrently in the organism. In natural environments, possible recontamination events, due to the reminiscence of toxic algal cells or the resuspension of particles from the sediment containing domoic acid may occur. Consequently, it explains why the depuration rates obtained in our study were slightly lower than those calculated from experimental studies, as discussed in Blanco et al. (2021).

Given the relatively low variability observed in depuration rates, we were able to investigate the potential factors influencing their magnitude. A comparison of depuration rates across provinces and years revealed no correlation with the tested variables: initial maximum concentration and environmental conditions (temperature, salinity and chlorophyll-*a*). No correlation (linear or nonlinear) between initial maximum domoic acid concentration and depuration rates could be observed, suggesting that higher toxin concentrations in tissues do not necessarily lead to faster depuration rates. However, during the investigation, we saw that estimations of depuration rates were more accurate (higher *R*^2^ value) when the initial domoic acid concentrations were high, often also associated with a higher number of points in the dataset for the contamination event. This is related to the methodology employed for the sampling process: when domoic acid concentration is above the regulatory threshold, samples are taken and analysed until the concentration returns to a level below the regulatory limit.

Furthermore, our results highlighted that depuration rates did not seem influenced by environmental conditions, at least for the ones tested in our study: temperature, salinity and chlorophyll-a concentration. Initially, we had hypothesised that depuration rates may be greater with higher seawater temperature, given its known effects on physiological processes, as well as with higher chlorophyll-*a* concentrations, as a proxy for food availability. We also considered salinity based on previous findings by Blanco et al. (2006), who reported a link between depuration rates and a covariation of both temperature and salinity, but found no significant relationship between depuration rates and food quantity. These outcomes are consistent with those reported by Blanco et al. (2006) and some results of Novaczek et al. (1992) on mussel contamination by domoic acid. In their study, Novaczek et al. (1992) examined the impact of temperature, salinity and food on the depuration rate of *M. edulis* and their findings revealed no effect of food, as found in the present study. Additionally, they observed a non-significant relationship with salinity, although a negative trend was noted, which is consistent with our findings, as indicated by a negative slope for the linear regression with salinity in spring and summer. However, their study revealed a positive correlation between depuration rate and temperature. These findings suggest that the link between depuration rates and individual bioenergetics remains unclear and warrants further investigation. Nevertheless, this may facilitate the definition of a unique depuration rate for this species and the identification of a method for predicting domoic acid depuration based on this common rate.

### A tool for predicting king scallop domoic acid depuration dynamics and concentrations

Predicting contamination events is crucial for effective fishery management, as the location and duration of closures influence the economic and operational impact on the activities (Chenouf et al., 2023). Given the unique environmental and regulatory characteristics of each province, mitigation strategies will vary; for example, closing specific zones, as performed in the Bay of Seine, or shifting efforts to other exploited species as previously observed in the Bay of Brest. In this context, our findings could contribute to the development of adaptive management strategies for professionals facing ASP events. Indeed, the ability of predicting with an anticipation of two or three months before the opening of the fishing season if the fishery will effectively open or not due to domoic acid contamination is crucial for professionals to reduce financial losses. While several studies have explored the forecasting of harmful algal blooms and toxin presence in bivalves (Bouquet et al., 2023, 2022), particularly in relation to *Pseudo-nitzschia* species and mussel contamination (Aláez et al., 2021), or harmful algal blooms forecasting using remote-sensing analyses (Zahir et al., 2024), to our knowledge, no research has focused on predicting the depuration dynamics of commercially exploited bivalves after a toxic bloom event. The main objective of this study was to evaluate the feasibility of predicting domoic acid concentrations in *P. maximus* tissues at the beginning of the fishing season based on domoic acid contamination levels measured in the spring after potential bloom of *Pseudo-nitzschia* spp. Using previous results discussed, we applied the median depuration rate of 0.0046 *d*^−1^ to each event, with an uncertainty range defined by the 25^*th*^ and 75^*th*^ percentiles (0.0033 *d*^−1^ and 0.0067 *d*^−1^). A prediction was considered accurate when at least 50% of the data points for a given event fell within this interval. Our method has proven effective, successfully predicting more than half of the events, with accuracy improving to 68% after refining the initial values and contamination event definitions in order to consider the end of the contamination period (*i.e*. when the bloom of toxic *Pseudo-nitzschia* is over) and events of recontamination. Thus, we recommend considering these insights when predicting new domoic acid concentration values.

As a second approach to predict domoic acid depuration, we explored forecasting toxin concentrations on a specific date. The aim was to determine an optimistic scenario that would allow to predict domoic acid concentrations close to the regulatory threshold while ensuring our predictions always overestimated the measured depuration. Indeed, predicting a reopening of the fishery activity before or after the effective one will not have the same consequences for professionals. In the event of the predicted date of reopening (*i.e*. the date at which concentration levels fall below the regulatory threshold) falling after the real authorised date, professionals may miss the fishing season if their vessel were not fitted out or they have not purchased a licence. In contrast, an early prediction of the opening date has less impact, and may lead to unnecessary fitting out of the boat without the authorisation to harvest scallops. Thus, we decided to predict the earliest reopening date for fishing in the event of contamination, considering it was the best advice to provide to management teams such as regional and departmental fishery committees, keeping in mind that it could be postponed. This was defined by the most optimistic predictive scenario defined in section 2.3.5. The variability around the initial concentration considered in this scenario, was defined based on observed inter-individual variability in individual domoic acid concentrations from (García-Corona et al., 2024). From an operational point of view, 81% of our predictions met the conditions for a satisfying result under this optimistic scenario. However, a key challenge was the limited availability of data on actual fishery reopening dates or cases when domoic acid concentrations were near the regulatory threshold, complicating direct comparisons between predictions and observations. Despite this limitation, our findings highlight the potential of integrating depuration modelling into decision-support tools, in order to assist fishery managers in making informed decisions ahead of the fishing season.

Our findings highlighted the critical importance of knowing with precision the initial domoic acid concentration. Indeed, the high variability in this value strongly influenced the outcomes of the predictions of the date at which domoic acid concentration is close to the regulatory threshold of 20 *mg kg*^−1^. We therefore recommend using the concentration measured at the end of the contamination event, corresponding to the end of the bloom, or the first concentration event after the bloom, to not bias the depuration rate, which corresponds to a balance between accumulation and depuration rates during the contamination process (Blanco et al., 2021). This was demonstrated by several cases, where considering the value at the end of the contamination period led to more accurate predictions, such as at the Gisement Le Fret (Intermediate Finistère) in 2017, with the adjustment of the initial value in July instead of May. Thus, for further fishery management purposes, considering sampling in summer and early autumn, following a *Pseudo-nitzschia* bloom in spring, could help refine the predictions. Furthermore, our results showed that the most reliable predictions were obtained when the initial domoic acid concentration was high and far from the regulatory limit of 20 *mg kg*^−1^. This is an inherent bias of the model: as concentrations decrease toward zero, small variations become more noticeable. Nevertheless, as demonstrated by our findings and previous studies (Husson et al., 2016; Blanco et al., 2021), the possibility of recontamination remains important and cannot be predicted with our approach. The only solution, for now, is to recalculate predictions when recontamination is detected through monitoring.

In order to assess the situations when the global dynamics of domoic acid depuration are difficult to predict, we highlighted and studied specific cases for which their estimated depuration rates were often at the extremes of the distribution (slow or rapid). This was particularly visible with the prediction of specific domoic acid concentrations for which 15 events were difficult to predict. Firstly, low estimated depuration rates in the first part of the study, might be linked to recontamination events, either due to the presence of toxic Pseudo-nitzschia spp. cells in the water column, which would need to be considered when estimating the depuration rates but cannot be predicted, and either due to a resuspension of domoic acid trapped into the sediment, as discussed in Husson et al. (2016); Blanco et al. (2021). Domoic acid contamination in bivalves and abundance of *Pseudo-nitzschia* are often linked but the difficulty remains in understanding the relationship between both variables (Belin et al., 2021; Husson et al., 2016) and sometimes the relationships are not clear (Giménez Papiol et al., 2013). Moreover not all species of *Pseudo-nitzschia* are toxic and the cell toxicity varies among species and strains (Bates et al., 2018; Sauvey et al., 2019). Additionally, it is known that inter-individual variability of domoic acid concentration is high for *P. maximus*, thus it can be difficult to distinguish between variability in concentration levels between two groups of individuals (10 individuals per group in the REPHYTOX sampling strategy) or a recontamination due to *Pseudo-nitzschia* cells in the water column or resuspension of cells from the sediment. On the other hand, rapid depuration, considered for rates above 0.01 *d*^−1^, was estimated for some events, such as at Quiberon-concessions in 2008 and at Golfe La Teignouse in 2013 (Morbihan), at Manche Est Large zone 15 (North Normandie) in 2012, and in the Pertuis and at the Gisement Roscanvel (Intermediate Finistère) in 2021. For the zone of Pertuis d’Antioche in 2021, we distinguished a high temperature value, based on the REPHY dataset, for the spring and summer seasons, higher than the ones of all other locations. Even if we did not find a correlation between depuration rates and temperatures, the question still remains open for very specific cases. For the others, the difficulty of predicting specific events could also be due to the uncertainty of the sampling method. In France, the sampling for the sanitary monitoring is carried out by fishermen, and not at a fixed spatial point with systematic sampling. Thus, there can be variations depending on the sampler; the individuals could be taken from different locations in one area or selected based on other criteria. This could lead to a difference in terms of domoic acid concentrations and thus impacting the depuration dynamic observed. On the other hand, the variation in domoic acid concentrations can be even more pronounced when considering large zones such as the ones defined in the Bay of Seine. Indeed, the GPS coordinates given with the samples for these locations showed that they were not sampled at the same place and could explain the high variability in concentrations observed. In this province, hydrodynamism has already been studied with modelling and observations and total production of diatoms and surface diatom concentrations in several months in 1978 have been mapped for this region (Cugier et al., 2005). This highlights the possibility of variation from one sampling point to another in terms of toxic cell concentration and thus in terms of bivalve contamination.

Finally, in addition to the predictive tool for domoic acid depuration dynamics, we assessed whether it was possible to rely on other shellfish as sentinel species, in the same area as *P. maximus*, to detect early contamination and/or assess the level of contamination in king scallops. To this end, we compared the domoic acid concentrations in other shellfish and *P. maximus* at the same locations but in the spring for other shellfish species and during the fishing season for *P. maximus*. The aim was to determine if knowing the level of contamination of other shellfish in the spring, when *P. maximus* is not exploited, could give insight into the potential contamination of *P. maximus* at the beginning of the fishing season (starting in October). Our results showed that whenever domoic acid is detected in another shellfish species (concentration above 1 *mg kg*^−1^), in 35% of cases king scallops are contaminated (domoic acid above the regulatory threshold of 20 *mg kg*^−1^) during the fishing season, this percentage increased up to 90% when considering detection of domoic acid in king scallops. If one shellfish species is contaminated with domoic acid above the regulatory threshold (20 *mg kg*^−1^), there is an 85% probability that king scallops will be contaminated during the fishing season. This suggests that detection of domoic acid in various species in the same areas is clearly correlated. However, we have seen that it is, for the moment, not possible to determine a correlation between the domoic acid concentrations in other bivalves and king scallops. Thus, it is not possible to estimate domoic acid concentrations in *P. maximus* when measuring the concentration in other bivalves. In our case, using a sentinel species to assess domoic acid contamination in king scallops is not an effective solution. This was already suggested by Belin et al. (2021) who discussed the fact that mussels were first used as sentinel species for diarrheic shellfish toxin detection, but it was not possible for paralytic and amnesic shellfish toxins using REPHY and REPHYTOX monitoring programmes, because king scallops depurate these toxins at a very low rates compared to other species. Similarly in Canadian coastlines, mussels were used as sentinel species for ASP and PSP because they represented the highest risk of contamination in this region (Rourke et al., 2021). However they also showed that depending on the toxin and the zone assessed, other species should also be monitored if they might represent a higher food safety risk, especially if the depuration rates are different, as for *P. maximus*. Hence, we recommend that upon the detection of domoic acid in a shellfish species during the spring months in an area where scallops are present, king scallops should be sampled during the summer months in order to assess the potential concentration in October, when the fishery activity starts.

### Solution for fisheries in France and extension to other countries

The fishery of king scallops in France remains open, in contrast to other countries such as Spain and Ireland. Nevertheless, the activity is confronted to prolonged periods of closure due to domoic acid contaminations. From this assessment, the MaSCoET (Maintaining the shellfish stock in the context of harmful algal blooms) scientific project was financed by the France Filière Pêche fishermen’s association with the objective of understanding the domoic acid contaminations and finding solutions for anticipating toxic *Pseudo-nitzschia* blooms and/or accelerating the decontamination of king scallops. One question posed by professionals during the course of this project concerned the feasibility of predicting domoic acid concentrations in king scallops at the opening of the fishing season in October. Indeed, the ability to determine during summer the risk of purchasing a licence, equipping a vessel, or whether it is preferable to engage in alternative activities is crucial for fishermen. It can contribute to reducing the economic losses incurred as a result of domoic acid contamination.

The strength of our study is the ability to analyse a large dataset of domoic acid concentrations in shellfish species along the French coasts from the operational monitoring REPHYTOX (REPHYTOX - French Monitoring program for phycotoxins in marine organisms, 2023), with an ultimate objective of offering an operational tool that is user-friendly and can be employed by fishery managers. Firstly, the comparison between species has enabled to propose a methodology to estimate the concentration of domoic acid in king scallops at a later date. We recommend that, when domoic acid is detected in one shellfish species during the spring period after a *Pseudo-nitzschia* bloom, king scallops should be sampled during summer for domoic acid quantification. This will enable fishing managers to anticipate contamination of king scallops three months before the opening of the fishing season. Secondly, the depuration model can be employed to predict domoic acid concentration in king scallop at a specified date (particularly in October, for the opening of the fishing season) based on the concentration at an earlier date (spring or summer after a contamination event). This does not consider potential recontamination, but it was found to be satisfactory for 81% of the events defined for France between 2004 and 2024. The development of an accessible online application for regional and departmental fishery committees is a future aim.

In light of the previous results and discussion, the implementation of management solutions could be considered, with the selection of such solutions being determined by the specific characteristics of the designated fishing zone. To illustrate this point, in the Bay of Seine, several zones can be defined with opening and closing dates depending on the domoic acid concentrations earlier in the year. In smaller king scallop beds, such as the Bay of Brest, the Bay of Morlaix or the Pertuis, potential management solution using our estimations can consist in choosing or not to purchase the fishing licence, to equip the vessel with specific dredging equipment, and in anticipating the fishing season several months in advance. This study was conducted on a French dataset; however, we trust that the method and the findings can be replicated in other regions having regular monitoring data such as Ireland and the Isle of Man. The efficacy of the managing solutions will be region-specific, including the type of assessment of domoic acid concentration in shellfish, as well as the organisation of fishery activity.

## 5. Conclusion

This study is one of the few to investigate depuration rates in a highly valuable bivalve species, the king scallop (*Pecten maximus*). By analysing 20 years of domoic acid concentration data from the in-situ sanitary monitoring programme REPHYTOX, we identified contamination events affecting the French Atlantic and English channel coastlines between 2004 and 2024. Estimating depuration rates and examining their spatio-temporal variability and correlations with the environmental conditions allowed us to define a common depuration rate applicable across different years and provinces.

Our findings demonstrate that a simple yet widely used statistical model (exponential decay) can effectively describe the majority of domoic acid contamination events recorded along the French coastline over the past two decades. This model provides a valuable predictive tool for fishery managers, helping to assess whether domoic acid concentrations in *P. maximus* tissues will fall below the regulatory threshold before the fishing season, particularly when contamination occurs in the spring. Looking ahead, our ultimate goal is to develop an accessible online application for regional and departmental fishery committees based on this model. This tool would enable managers to input domoic acid concentrations observed in the spring and receive an estimate of potential fishery closures under the most optimistic scenario.

To further refine predictions and better account for inter-event variability, we propose exploring more complex models, such as the mechanistic bioenergetic approach. Such models could help to represent inter-individual variability, improving accuracy and providing a range of predictions based on inter-individual variability, well known for domoic acid retention in pectinid species. Similar models have already been developed, applied to other harmful algal blooms, such as *Alexandrium minutum*, and its paralytic shellfish toxins in oysters (Pousse et al., 2019).

This study provides new insight into domoic acid depuration dynamics in king scallops and, by advancing predictive modelling approaches, this research contributes to more effective fishery management strategies, reducing economic impacts while ensuring food safety.

## 6. Acknowledgements

The authors would like to thank the members of the Ifremer coastal laboratories and all other persons who performed the sampling and analyses, as well as the national coordination team of REPHY-REPHYTOX, those who created and maintained the Quadrige database, and all experts who contributed to these datasets. This work received financial support from the research project “MaSCoET” (Maintien du stock de coquillages en lien avec la problématique des efflorescences toxiques) financed by France Filière Pêche and Brest Métropole. Eline Le Moan was recipient of a doctorate fellowship financed by France Filière Pêche and Région Bretagne.

---

**Table S1:** Coordinates (WGS84) of the provinces defined in this study, with minimum longitude and latitude (in decimal degrees) corresponding to the southwest corner of the square and maximum longitude and latitude corresponding to the northeast corner of the square for each province.

| Province | Minimum longitude | Maximum longitude | Minimum latitude | Maximum latitude |
| --- | --- | --- | --- | --- |
| North | -0.3 | 2.1 | 50.4 | 51.8 |
| North Normandie | 0.25 | 1.8 | 49.6 | 50.4 |
| Bay of Seine | -1.4 | 0.3 | 49.1 | 50.1 |
| Western Normandie | -2.1 | -1.4 | 48.6 | 49.8 |
| Côtes d'Armor | -3.6 | -2.1 | 48.4 | 48.9 |
| North Brittany open sea | -5.3 | -2.1 | 48.9 | 50.0 |
| North Finistère | -4.8 | -3.6 | 48.5 | 48.9 |
| Intermediate Finistère | -5.3 | -4.2 | 48 | 48.5 |
| South Finistère | -4.9 | -3.6 | 47.6 | 48.0 |
| Morbihan | -3.6 | -2.4 | 47 | 47.8 |
| Loire | -2.4 | -1.6 | 46.4 | 47.4 |
| Pertuis | -1.7 | -1 | 45.6 | 46.4 |
| South west | -2.2 | -0.8 | 43.3 | 45.6 |

## References

Aláez, F.M.B., Palenzuela, J.M.T., Spyrakos, E., Vilas, L.G., 2021. Machine Learning Methods Applied to the Prediction of Pseudo-nitzschia spp. Blooms in the Galician Rias Baixas (NW Spain). ISPRS International Journal of Geo-Information 10, 199. doi:10.3390/ijgi10040199.

Álvarez, O.A., Rengel, J., Araya, M., Álvarez, F., Pino, R., Uribe, E., Díaz, P.A., Rossignoli, A.E., López-Rivera, A., Blanco, J., 2020. Rapid Domoic Acid Depuration in the Scallop Argopecten purpuratus and Its Transfer from the Digestive Gland to Other Organs. Toxins 12, 698. doi:10.3390/toxins12110698.

Bates, S.S., Bird, C.J., Boyd, R.K., Hanic, L.A., Jamieson, W.O., McCul-loch, A.W., Odense, P., Quiliiom, M.A., Sim, P.G., Thiboult, P., 1988. Investigations on the Source of Domoic Acid Responsible for the Outbreak of Amnesic Shellfish Poisonning (ASP) in Eastern Prince Edward Island. Technical Report 57. National Research Council.

Bates, S.S., Hubbard, K.A., Lundholm, N., Montresor, M., Leaw, C.P., 2018. Pseudo-Nitzschia, Nitzschia, and domoic acid: New research since 2011. Harmful Algae 79, 3–43. doi:10.1016/j.hal.2018.06.001.

Belin, C., Soudant, D., Amzil, Z., 2021. Three decades of data on phytoplankton and phycotoxins on the French coast: Lessons from REPHY and RE-PHYTOX. Harmful Algae 102, 101733. doi:10.1016/j.hal.2019.101733.

Blanco, J., Acosta, C., Bermúdez de la Puente, M., Salgado, C., 2002. Depuration and anatomical distribution of the amnesic shellfish poisoning (ASP) toxin domoic acid in the king scallop Pecten maximus. Aquatic Toxicology 60, 111–121. doi:10.1016/S0166-445X(01)00274-0.

Blanco, J., Acosta, C., Mari\∼no, C., Mu\∼niz, S., Mart\’in, H., Moro\∼no, \., Correa, J., Arévalo, F., Salgado, C., 2006. Depuration of domoic acid from different body compartments of the king scallop Pecten maximus grown in raft culture and natural bed. Aquatic Living Resources 19. doi:10.1051/alr:2006026.

Blanco, J., Moroño, Á., Arévalo, F., Correa, J., Salgado, C., Rossignoli, A.E., Lamas, J.P., 2021. Twenty-Five Years of Domoic Acid Monitoring in Galicia (NW Spain): Spatial, Temporal and Interspecific Variations. Toxins 13, 756. doi:10.3390/toxins13110756.

Blanco, J., Moroño, A., Franco, J., Reyero, M.I., 1997. PSP detoxification kinetics in the mussel Mytilus galloprovincialis>/i>. One- and two-compartment models and the effect of some environmental variables. Marine Ecology Progress Series 158, 165–175. doi:10.3354/meps158165.

Bouquet, A., Laabir, M., Rolland, J.L., Chomérat, N., Reynes, C., Sabatier, R., Felix, C., Berteau, T., Chiantella, C., Abadie, E., 2022. Prediction of Alexandrium and Dinophysis algal blooms and shellfish contamination in French Mediterranean Lagoons using decision trees and linear regression: A result of 10 years of sanitary monitoring. Harmful Algae 115, 102234. doi:10.1016/j.hal.2022.102234.

Bouquet, A., Thébault, A., Arnich, N., Foucault, E., Caillard, E., Gianaroli, C., Bellamy, E., Rolland, J.L., Laabir, M., Abadie, E., 2023. Modelling spatiotemporal distributions of Vulcanodinium rugosum and pinnatoxin G in French Mediterranean lagoons: Application to human health risk characterisation. Harmful Algae 129, 102500. doi:10.1016/j.hal.2023.102500.

Chenouf, S., Pérez Agúndez, J.A., Raux, P., 2023. Analysing the Socioeconomic Impacts of Fishing Closures Due to Toxic Algal Blooms: Application of the Vulnerability Framework to the Case of the Scallop Fishery in the Eastern English Channel. Sustainability 15, 12379. doi:10.3390/su151612379.

Commission, E., 2002. Commission Decision of 15 March 2002 establishing special health checks for the harvesting and processing of certain bivalve molluscs with a level of amnesic shellfish poison (ASP) exceeding the limit laid down by Council Directive 91/492/EEC. Official Journal of the European Communities, 65–66.

Cugier, P., Ménesguen, A., Guillaud, J.F., 2005. Three-dimensional (3D) ecological modelling of the Bay of Seine (English Channel, France). Journal of Sea Research 54, 104–124. doi:10.1016/j.seares.2005.02.009.

Douglas, D.J., Kenchington, E.R., Bird, C.J., Pocklington, R., Bradford, B., Silvert, W., 1997. Accumulation of domoic acid by the sea scallop (Placopecten magellanicus) fed cultured cells of toxic Pseudo-nitzschia multiseries. Canadian Journal of Fisheries and Aquatic Sciences 54, 907–913. doi:10.1139/f96-333.

FAO, 2025a. Gloabl Aquaculture Production. https://www.fao.org/fish-ery/en/collection/aquaculture?lang=en.

FAO, 2025b. Global Capture Production. https://www.fao.org/fish-ery/en/collection/capture?lang=en.

FranceAgriMer, 2022. Observatoire de La Formation Des Prix et Des Marges Des Produits Alimentaires. Technical Report.

García-Corona, J.L., Fabioux, C., Vanmaldergem, J., Petek, S., Derrien, A., Terre-Terrillon, A., Bressolier, L., Breton, F., Hegaret, H., 2024. The amnesic shellfish poisoning toxin, domoic acid: The tattoo of the king scallop Pecten maximus. Harmful Algae 133, 102607. doi:10.1016/j.hal.2024.102607.

Giménez Papiol, G., Casanova, A., Fernández-Tejedor, M., de la Iglesia, P., Diogène, J., 2013. Management of domoic acid monitoring in shellfish from the Catalan coast. Environmental Monitoring and Assessment 185, 6653–6666. doi:10.1007/s10661-012-3054-6.

Guillotreau, P., Bihan, V.L., Morineau, B., Pardo, S., 2021. The vulnerability of shellfish farmers to HAB events: An optimal matching analysis of closure decrees. Harmful Algae 101, 101968. doi:10.1016/j.hal.2020.101968.

Guiry, M., Guiry, G., 2021. AlgaeBase. World-wide electronic publication. https://www.algaebase.org.

Holland, D.S., Leonard, J., 2020. Is a delay a disaster? economic impacts of the delay of the california dungeness crab fishery due to a harmful algal bloom. Harmful Algae 98, 101904. doi:10.1016/j.hal.2020.101904.

Husson, B., Hernández-Fariñas, T., Le Gendre, R., Schapira, M., Chapelle, A., 2016. Two decades of Pseudo-nitzschia spp. blooms and king scallop (Pecten maximus) contamination by domoic acid along the French Atlantic and English Channel coasts: Seasonal dynamics, spatial hetero-geneity and interannual variability. Harmful Algae 51, 26–39. doi:10.1016/j.hal.2015.10.017.

Le Moan, E., Derrien, A., Fabioux, C., Jean, F., Lassudrie, M., Terre-Terrillon, A., Hégaret, H., Flye-Sainte-Marie, J., 2025. Domoic acid depuration rates in king scallops, analyses across French provinces from 2004 to 2024. doi:10.17882/106299.

Lelong, A., Hégaret, H., Soudant, P., Bates, S.S., 2012. Pseudo-Nitzschia (Bacillariophyceae) species, domoic acid and amnesic shellfish poisoning: Revisiting previous paradigms. Phycologia 51, 168–216. doi:10.2216/11-37.1.

Lundholm, N., Bernard, C., Churro, C., Escalera, L., Hoppenrath, M., Iwataki, M., Larsen, J., Mertens, K., Moestrop, \., Murray, Salas R., Tillman, U., Zingone, A., 2009. IOC-UNESCO Taxonomic Reference List of Harmful Micro Algae. https://www.marine-species.org/hab/aphia.php?p=taxdetails&id=149151.

MacDonald, B.A., Bricelj, V.M., Shumway, S.E., 2016. Chapter 7 - Physiology: Energy Acquisition and Utilisation, in: Shumway, S.E., Parsons, G.J. (Eds.), Developments in Aquaculture and Fisheries Science. Elsevier. volume 40 of Scallops, pp. 301–353. doi:10.1016/B978-0-444-62710-0.00007-9.

Mafra, L.L., Bricelj, V.M., Fennel, K., 2010. Domoic acid uptake and elimination kinetics in oysters and mussels in relation to body size and anatomical distribution of toxin. Aquatic Toxicology 100, 17–29. doi:10.1016/j.aquatox.2010.07.002.

Novaczek, I., Madhyastha, M.S., Ablett, R.F., Donald, A., Johnson, G., Nijjar, M.S., Sims, D.E., 1992. Depuration of Domoic Acid from Live Blue Mussels (Mytilus edulis). Canadian Journal of Fisheries and Aquatic Sciences 49, 312–318. doi:10.1139/f92-035.

Pousse, E., Flye-Sainte-Marie, J., Alunno-Bruscia, M., Hégaret, H., Rannou, E., Pecquerie, L., Marques, G.M., Thomas, Y., Castrec, J., Fabioux, C., Long, M., Lassudrie, M., Hermabessiere, L., Amzil, Z., Soudant, P., Jean, F., 2019. Modelling paralytic shellfish toxins (PST) accumulation in Crassostrea gigas by using Dynamic Energy Budgets (DEB). Journal of Sea Research 143, 152–164. doi:10.1016/j.seares.2018.09.002.

Pulido, O.M., 2008. Domoic Acid Toxicologic Pathology: A Review. Marine Drugs 6, 180–219. doi:10.3390/md6020180.

R core team, 2022. R: A language and environment for statistical computing. R foundation for Statistical computing.

REPHY - French Observation and Monitoring program for phytoplankton and hydrology in coastal waters, 2023. REPHY dataset. French Observation and Monitoring program for Phytoplankton and Hydrology in coastal waters. Metropolitan data. doi:10.17882/47248.

REPHYTOX - French Monitoring program for phycotoxins in marine organisms, 2023. REPHYTOX dataset. French Monitoring program for Phycotoxins in marine organisms. Data since 1987. doi:10.17882/47251.

Rourke, W.A., Justason, A., Martin, J.L., Murphy, C.J., 2021. Shellfish Toxin Uptake and Depuration in Multiple Atlantic Canadian Molluscan Species: Application to Selection of Sentinel Species in Monitoring Programs. Toxins 13, 168. doi:10.3390/toxins13020168.

Sauvey, A., Claquin, P., Le Roy, B., Le Gac, M., Fauchot, J., 2019. Differential Influence of Life Cycle on Growth and Toxin Production of three Pseudo-nitzschia Species (Bacillariophyceae). Journal of Phycology 55, 1126–1139. doi:10.1111/jpy.12898.

Sellner, K.G., Doucette, G.J., Kirkpatrick, G.J., 2003. Harmful algal blooms: Causes, impacts and detection. Journal of Industrial Microbiology and Biotechnology 30, 383–406. doi:10.1007/s10295-003-0074-9.

Wohlgeschaffen, G.D., Mann, K.H., Subba Rao, D.V., Pocklington, R., 1992. Dynamics of the phycotoxin domoic acid: Accumulation and excretion in two commercially important bivalves. Journal of Applied Phycology 4, 297–310. doi:10.1007/BF02185786.

Wright, J.L.C., Boyd, R.K., de Freitas, A.S.W., Falk, M., Foxall, R.A., Jamieson, W.D., Laycock, M.V., McCulloch, A.W., McInnes, A.G., Odense, P., Pathak, V.P., Quilliam, M.A., Ragan, M.A., Sim, P.G., Thibault, P., Walter, J.A., Gilgan, M., Richard, D.J.A., Dewar, D., 1989. Identification of domoic acid, a neuroexcitatory amino acid, in toxic mussels from eastern Prince Edward Island. Canadian Journal of Chemistry 67, 481–490. doi:10.1139/v89-075.

Zahir, M., Su, Y., Shahzad, M.I., Ayub, G., Rehman, S.U., Ijaz, J., 2024. A review on monitoring, forecasting, and early warning of harmful algal bloom. Aquaculture, 741351 doi:10.1016/j.aquaculture.2024.741351.

